# High-throughput microfluidic encapsulation of cells in extracellular matrix microbeads for microtissue engineering

**DOI:** 10.64898/2026.09.30.755758

**Authors:** Debjita Mukherjee, Magdalena Radecka-Rudzinska, Konrad Giżyński, Marta Klak, Michal Wszola, Jan Guzowski

**Affiliations:** Institute of Physical Chemistry, Polish Academy of Sciences, Marcina Kasprzaka 44/52, 01-224, Warsaw, Poland; Foundation of Research and Science Development, Al. Prymasa Tysiąclecia 79A, 01-242 Warsaw, Poland

## Abstract

Microfluidics-assisted encapsulation of cells in hydrogel microbeads has found diverse applications in tissue engineering, disease modelling, and drug testing. However, upscaled production of submillimeter-sized microbeads, each encapsulating optimally tens to hundreds of living cells, remains a challenge. Excessive viscous stresses or non-homogeneous crosslinking reactions hamper droplet monodispersity, hydrogel integrity, and/or cell viability. Herein, we present a device consisting of (i) tens of parallel step-emulsifiers arranged radially around the hydrogel inlet and (ii) an in-flow UV crosslinking system capable of continuous generation and crosslinking of up to around 45000 microbeads/min, each encapsulating *N*∼50 cells, at cell viability exceeding 80%. We use gelatin methacryloyl (GelMA) as the biomaterial of choice and establish a unique “sweet spot” in terms of UV wavelength, irradiance, and exposure time—the latter controlled via the net applied flow rate and the length of the UV-exposed microchannel—at which we achieve an optimal balance between long-term microbead stability and cell viability. In particular, we observe a strong correlation between reduced cell viability and poor crosslinking efficiency, the latter attributed to the presence of a spontaneous core-shell structure. We further study the impact of added methacrylated hyaluronic acid (HAMA) and extracellular matrix (ECM) on cell proliferation and microtissue formation in the case of triple-negative breast cancer (TNBC) cells for future application in drug testing. The upscaled production of the encapsulated microtissues demonstrated here provides new perspectives for both drug testing and cell therapies where the precision in microenvironment formulation as well as production throughput remain of primary importance.

## Introduction

There has been an increasing interest in reproducible fabrication of microtissues, that is, microscopic cell aggregates mimicking some of the native tissue functions, for applications in tissue engineering, drug screening, or cell therapies^1–3^. Droplet microfluidics appears as a particularly promising approach in this matter by supporting high-throughput generation of cell-laden microdroplets as highly monodisperse microcarriers^4–6^. Apart from providing efficient compartmentalization, the droplets may also co-encapsulate dissolved biopolymers, which, after crosslinking, provide a soft-solid hydrogel matrix, closely resembling the native extracellular matrix. In comparison to more conventional methods of microtissue formulation--such as cell self-aggregation in microwells^7^ -- the microfluidic approach offers (i) significantly limited biomaterial consumption, (ii) superior monodispersity of the generated microtissues^8^, and (iii) enhanced throughput of formulation. Importantly, it also avoids time-consuming and labour-intensive experimental procedures^9^. Overall, by consistently providing a well-controlled cell microenvironment, droplet microfluidics opens the way to systematic studies on cell responses under various external cues at the microtissue-level^10^, which remain crucial, e.g., in applications such as cell therapies, drug screening, or stem-cell niche engineering^11–13^. Future prospects include widespread adoption of such miniaturized human tissue models as a replacement for animal models in line with 3R/6R guidelines^14^.

Despite significant progress in this field, several issues remain unresolved. One of the crucial challenges is elevating the throughput of microtissue generation. Currently, the main limitation is not the formation of the aqueous or hydrogel microdroplets per se, which, with the available methods, can easily reach litres per hour^15–17^, but rather the specific requirements associated with cell viability, proliferation, aggregation, and tissue maturation. To this end, the generated microbeads should optimally

i. encapsulate a sufficiently high number of cells in each droplet, with a reasonable minimal number N ∼ 50, which at a typical cell concentration ∼ 10^7^ cells/mL corresponds to the droplet diameter roughly *D* ∼ 200 um;
ii. consist of a biopolymer matrix closely resembling the extracellular matrix (ECM) encountered in the native tissue;
iii. be generated under mild physico-chemical conditions, that is, under limited viscous stresses and mild biopolymer crosslinking reactions (e.g., limited exposure to UV and/or cytotoxic chemical reactions);
iv. remain stable (non-degradable) under long-term culture (3-7 days; in some cases, even several weeks)—to provide a physical barrier against the external environment, allow tissue maturation, and facilitate further downstream manipulation and analysis.

Despite a significant body of literature reporting microfluidic formulation of hydrogel microbeads for microtissue engineering, the currently available methods often violate some of the above-mentioned requirements or otherwise suffer from a limited throughput.

Many of the available techniques reporting high-throughput microgel generation have been optimized for single-cell encapsulation^18,19^. These high-throughput methods have been used for single-cell sorting and screening, such as for rapid downstream analysis, such as DNA/RNA sequencing^20–22^. However, the microgels generated using these techniques are exceedingly small and therefore cannot support multi-cell aggregation. In fact, multi-cell encapsulation is absolutely necessary to rapidly provide both direct and indirect cell-cell interactions, which impact their phenotype and/or metabolism^23^.

Microtissue formation requires both cell-cell as well as cell-ECM interactions, where the latter is generally modulated by the amino acid sequences, like RGD (amino acids R: Arginine, G: Glycine, and D: Aspartic acid), present in the hydrogel used for encapsulation^24^. The earlier reported ultra-high-throughput multi-cell encapsulation methods, such as in-air microfluidics^25^, were typically based on ionic alginate crosslinking. Even though the in-air systems provide uniquely high throughputs, maintaining monodispersity of the cell-laden droplets remains a challenge in such systems because the presence of cells seems to interfere with the droplet pinch-off process. Other factors such as droplet impact in the ionic bath during generation may also significantly affect droplet shape^26^. Admittedly, the use of alginate as the hydrogel of choice has several benefits, like consistent physico-chemical properties and ultra-fast crosslinking, but the native alginate lacks cell-binding peptides or natural components of the ECM^27^ and therefore cannot support proper cell and tissue phenotype. In comparison, natural protein-based hydrogels such as gelatin, fibrin, and collagen naturally support cell adhesion and proliferation (and in some cases also differentiation), and are inherently viscoelastic and capable of faster stress relaxation,^28^ unlike hydrogels like alginate and PEG ^29^, and therefore are much more promising as ECM-mimics^30^. The use of tissue-specific decellularized ECMs (dECM) further improves cell viability and proliferation, and may provide a highly specific microenvironment resulting in decreased immunogenic responses, essential, e.g., in transplantation-based treatments^31^. Even though microdroplets can be generated from dECMs alone, their structural stability is temperature-dependent^32,33^. To stabilize dECM-based microspheres, various approaches have been employed, such as methacrylation of dECM^34^, adding coatings to the generated microspheres,^35^ or further addition of chemically crosslinkable hydrogels such as GelMA, alginate, PLGA, etc. ^36–39^ Such biomaterial combinations can be used to generate accurate microenvironments for tumor models and regenerative therapies, with microfluidics aiding in the formation of structurally advanced capsules such as core-shell structures, hemispheres, or other structures^6,40–42^.

Several methods have been developed to crosslink and stabilize individual hydrogel beads, including physical and chemical crosslinking, enzymatic linkage, and/or photo-crosslinking using UV light^43^. Chemically-driven covalent crosslinking using crosslinkers like glutaraldehyde, or catalyst-free methods with heavily modified polymeric precursors, or residual monomers from hydrogels like polyacrylamide (acrylamide monomer is a neurotoxin/carcinogen), often results in cytotoxicity^44–46^. Gelatin-derived hydrogels, on the other hand, can be easily modified to allow photocrosslinking by free-radical polymerization in the presence of a photo-initiator^47^. In particular, gelatin methacryloyl (GelMA) hydrogels facilitate fast fabrication of the microbeads via photocrosslinking of the pre-generated GelMA droplets, while also providing a soft hydrogel matrix, highly favourable for cell growth and proliferation. Another added value of GelMA is the amenability to local enzymatic degradation by the cells, which allows for recapitulating physiological tissue formation processes driven by cell migration^48^. One has to remember that photocrosslinking is also prone to causing cytotoxicity and needs careful monitoring and optimization of the associated factors, such as UV light wavelength, irradiance, choice/concentration of photoinitiator, and exposure time, to avoid cell viability issues^49,50^. We provide such optimization in the current work in the case of droplets generated at high-throughput using microfluidics.

In particular, we demonstrate a high-throughput hydrogel microbead production system exploiting multiple parallel droplet generators and ‘on-the-fly’ photo-crosslinking of the hydrogel-precursor microdroplets using an external LED panel as a light source in the UV-Vis range (385 nm - 405 nm). The system integrates two chips connected via tubing: the droplet-generation chip (D-chip), consisting of N = 64 equivalent step-emulsification junctions producing the droplets at ultra-high throughput (>10^4^ droplets /min), and the droplet-crosslinking chip (C-chip), consisting of a meandering channel placed in front of the LED panel, thus warranting direct and homogeneous high-intensity illumination. The LED panel, together with the C-chip, is additionally enclosed in a non-transparent crosslinking chamber to eliminate any possible exposure of the D-chip and thus prevent hydrogel crosslinking, which otherwise could lead to clogging of the droplet generators.

In this work, we carefully examine a variety of factors affecting microbead crosslinking efficiency and cell viability at retained high throughput of the microbead generation. In particular, we find that the generation of well-crosslinked (long-term stable) microbeads with encapsulated viable cells can be achieved by: (i) matching the illumination wavelength with the photoinitiator absorption band, (ii) the use of an optimized UV irradiance (not too high, not too low), (iii) sufficiently long exposure time (> 10 seconds in our case), (iv) providing thermal stability, e.g., via setting a sufficiently large distance between the C-chip and the light source to avoid droplet overheating. Under carefully fine-tuned conditions, we demonstrate generation and crosslinking of GelMA microbeads of diameter *D*∼200 um, each encapsulating around *N*∼50 cells, at a throughput of up to 45000 beads/min. The process retains high monodispersity of the beads (CV < 5%), high homogeneity in cell distribution per bead (CV < 30%), long-term stability of the beads (up to 14 days in culture), and the viability of the encapsulated cells at >80% (24 h post-encapsulation in the case of model fibroblasts and cancer cells).

Finally, we also tested the addition of HAMA and decellularized porcine ECM to the droplets along with GelMA. We found that the composite GelMA-HAMA-ECM hydrogel supports better viability and proliferation rates of the encapsulated TNBC cells as compared to the pure GelMA or GelMA-HAMA mixture, resulting in mature breast microtumors already at Day 5 of culture. This demonstrates potential for application of the system in future drug-screening studies. Owing to the native pancreatic ECM component, we also foresee possible applications of our high-throughput system in beta cell encapsulation and islet engineering, as well as in other cell therapy applications where the upscaled production remains a bottleneck.

## 2. Materials and Methods

### 2.1. Microfluidic chips fabrication and design

The channels of the microfluidic chips were designed using AutoCAD software and micromilled in polymethylmethacrylate (PMMA). Polydimethylsiloxane (PDMS) (SYLGARD™ 184 Silicone Elastomer Kit, Dow Corning Corporation, Michigan, United States) was used to fabricate the microfluidic chips. PDMS pre-polymer was mixed with the curing agent in a 9:1 ratio, thoroughly degassed in a vacuum pump, poured onto the PMMA chip, and cured at 80°C for 12 hours to produce a negative PDMS master mold.

Next, the PDMS master was silanized with trichloro(1H,1H,2H,2H-perfluorooctyl) silane (PFOCTS; Gelest Inc., United States). Another batch of the same PDMS pre-polymer with the curing agent was poured onto the PDMS master, carefully degassed to remove all air bubbles, and left at 80°C for 4-6 hours. Then, the positive copy of the PDMS chip was removed from the mold and bonded to a glass slide after plasma treatment for 2 minutes. This procedure was followed for both the droplet-producing chip (D-Chip, Figure 1.e.) and the crosslinking chip (C-Chip; Figure 1.g.).

**Figure 1.**
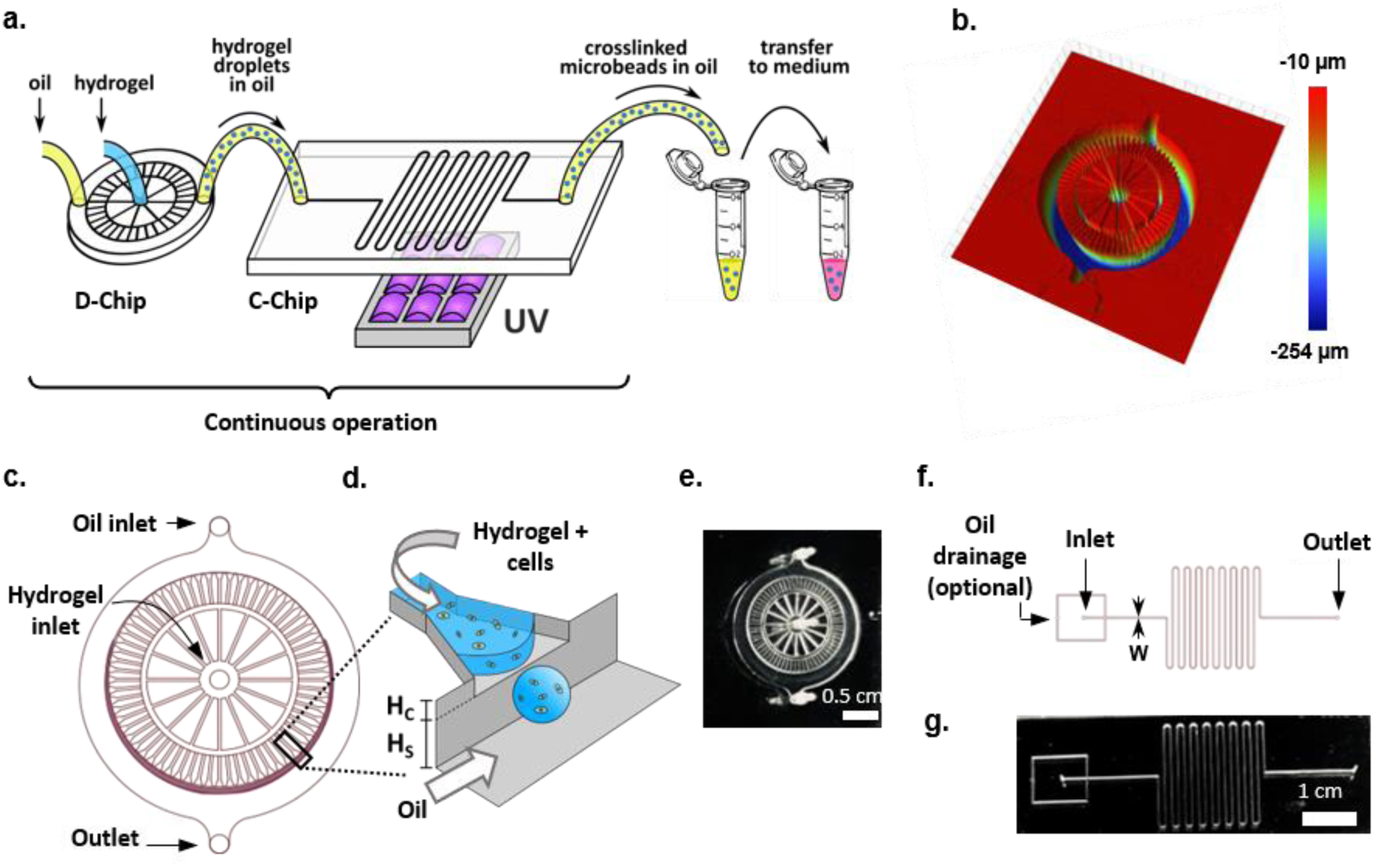
System components and design. **a.** Schematic of the complete system consisting of the droplet production chip (D-chip) forming hydrogel droplets in oil which are carried over to the crosslinking chip (C-chip) where the droplets crosslink as they pass before a UV LED lamp and are collected in oil and further washed and transferred to cell culture media. **b.** 3-D profilometric mapping of the D-chip. **c.** Layout of the droplet generator chip (D-chip). **d.** The droplet generation process from a single channel by step-emulsification. **e.** Image of the original D-chip fabricated in PDMS. **f.** Layout of the droplet crosslinking chip (C-chip); g. Image of the C-chip fabricated in PDMS.

The D-Chip consisted of an inlet for the droplet phase, splitting the flow into 16 radial channels (*W* = 120 μm, *H* = 150 μm) terminating at a concentric circular channel acting as a pressure reservoir (*W* = 250 μm, *H* = 150 μm). The circular channel supplied 64 shallow radial channels (*W* = 120 μm, *H* = 40 μm) serving as droplet generators and terminating at a circular edge of a droplet-collection channel (*H* = 200 μm) (Figure 1.b. and c.). In the following, for convenience, we denote by *H*_C_ the height of the shallow channels and by *H*_S_ the depth of the ‘step’ such that *H*_C_ + *H*_S_ equals the height of the droplet collection channel (Figure 1.d.).

The droplets formed spontaneously at the edge of the chamber via the process called step-emulsification ^51^. The wedge-shaped outlets of the droplet-generating channels gradually widened to facilitate the formation of the droplets and their pinch-off at the water-oil interface^52^. The flow of the continuous oil phase was supplied to the collection chamber via an inlet on one side, and the generated emulsion was collected at the outlet on the other side. The chamber widened towards the outlet to accommodate the increasing volume fraction of the droplets generated along the edge.

The C-Chip consisted of a meandering channel of square cross-section, *W* x *H* = 300 μm x 300 μm (Figure 1.f.). Two versions were used: Chip-A with a channel of total length *L* = 32.7 cm with 13 turns, or Chip-B with a total length *L* = 98.1 cm with 39 turns (Figure S1.d.). In both cases, the channels occupied the XY-footprint of 2.4 cm x 2.4 cm, consistent with the lateral dimensions of the LED array of the UV lamp. As the droplets flowed through the meandering channel, they were crosslinked gradually by free radical polymerization under UV exposure (Figure 1.a.). The depths of the channels in the micromilled devices were measured with an optical profilometer (Contour GT-G, Bruker, Massachusetts, United States) based on white light interferometry (Figure 1.b.). The profilometer used the software Vision 64 to acquire the images and calculate the dimensions of the channels.

#### 2.1.1. Numerical simulation of the two-phase microflows at the D-chip

A three-dimensional model of the D-chip channels was developed in COMSOL Multiphysics using the Two-Phase Flow, Level-Set interface. To reduce computational cost, a symmetry plane was introduced at *x* = 0.

The hydrogel phase (Fluid 1) was defined with a dynamic viscosity of *η*_hydrogel_ = 0.01 Pa ⋅ s and density *ρ*_hydrogel_ = 1000 kg/m^3^. The oil phase (Fluid 2) was characterized by *η*_oil_ = 0.0012 Pa ⋅ s and *ρ*_oil_ = 1614 kg/m^3^. Surface tension between the two phases was set to 5 × 10^−3^ N/m. At the inlets, constant volumetric flow rates were imposed. The hydrogel inlet flow rate was set to:

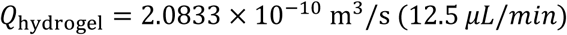

while the oil inlet flow rate was:

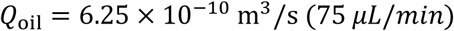

All remaining boundaries were defined as wetted walls with a contact angle of 1° (corresponding to the hydrogel phase Young’s contact angle of 179°) and a Navier slip length of 2 × 10^−6^ m. Gravity was included in the Navier–Stokes equations. The computational mesh was generated using the physics-controlled meshing option with the “Finer” setting. In the Level-Set formulation, the reinitialization parameter was set to 18.7 × 10^−3^ m/s, corresponding approximately to the maximum fluid velocity observed in the system. The parameter controlling the interface thickness was defined as 0.5 × ls. ep_default.

### 2.2. Single-run droplet generation and photo-crosslinking

A Cetoni neMESYS pump (Cetoni GmbH, Germany) and glass Hamilton syringes (1 mL for the hydrogel phase and 5 mL for the oil phase) were used to supply the flows. The hydrogel phase was loaded into a PE (polyethylene) tubing (inner diameter ID = 0.76 mm and outer diameter OD = 1.2 mm; Instech Laboratories Inc., USA), while the syringe was loaded with the oil phase to push out the hydrogel from the tubing (Figure S1.a.). This was done to significantly reduce issues regarding gravity-driven cell settling over time. The oil phase was a mixture of hydrofluoroether (Novec™ 7500, 3M, United States) with a fluorosurfactant (PFPE-PEG-PFPE; Chemipan R&D Laboratories, Poland) at 2% (w/w) concentration. The pump was connected to the D-chip with the PE tubing, where the droplets were produced. The process of microparticle generation was monitored with a stereomicroscope (Leica Microsystems, Germany).

The C-chip was mounted on a holder placed before the UV LED lamp fixed on a rail in a black PMMA box impervious to external light to allow accurate UV exposure to the droplets (Figure S3.e.). The C-chip was connected to the D-chip using PE tubing. The box was fitted with a UV-protectant screen to allow visual inspection of the crosslinking process (Figure S1.b.). A fan was fitted to the box wall to dissipate the heat generated by the LEDs. The crosslinked droplets were finally collected via an outlet tubing from the C-chip and washed to remove the oil phase. The schematic of the complete crosslinked droplet generation process can be seen in Figure 1.a.

The washing process was performed as follows (Figure S1.e.): After collection of the crosslinked emulsion droplets, i.e., the suspension of microbeads in oil in an Eppendorf tube, the suspension spontaneously separated (by gravity) into an oil-rich layer underneath the microbead-rich layer. Next, the oil layer was aspirated and discarded, and 500 µl of a 4:1 mixture of Novec 7500 and 1H,1H,2H,2H-perfluorooctanol (PFO; ThermoFisher Scientific GmbH, Germany) was added to the microbeads to neutralize the surfactant in the remaining residual oil. The suspension was mixed thoroughly via repeated pipetting. After the beads clustered at the top of the tube, the Novec-PFO mixture was removed from underneath the beads and replaced with pure Novec 7500. Washing with pure Novec was repeated three times. Finally, Novec was removed and replaced with 1 mL phosphate-buffered saline (PBS; PluriSelect Life Science, Germany) and centrifuged at 1200 rpm for 5 minutes. After centrifugation, three distinct layers were visible: PBS on top, the microbeads in the middle, and oil residue at the bottom. The oil layer was again carefully removed, and the PBS washing process was repeated thrice. After the final wash, PBS was replaced with fresh cell culture media, transferred to cell culture dishes, and cultured at 5 % CO_2_ and 37°C in a humidified incubator.

### 2.3. Characterization of the light sources

The custom-made UV-violet light (405 nm) LED lamp (further referred to as homemade lamp or HL) consisted of a square array of 5 x 5 = 25 LEDs at 5 mm spacing, where each LED had the output power P = 1W (Figure S1.c.i.). The lamp provided illumination intensities in the range 0-300 mW/cm^2^ as measured by an optical power meter (Figure S3.a.).

We used two LED lamps providing different wavelengths (385 nm and 405 nm) obtained from LucBei, China. Each lamp consisted of a square array of LEDs with a lateral footprint of around 2.5 cm x 2.5 cm and provided irradiance in the range 0-1600 mW/cm^2^ as measured by an optical power meter PM 100D (Thorlabs, Inc., United States) (Figure S3.a.).

To measure irradiance, the meter was attached to the holder, placed above the center of the LED array, and the distance between the light sensor and the LED lamp was varied in the range 0.5-5 cm (Figure S3.b.). The input amperage of the LED lamp was varied in the range 0.1 A - 3 A at different distances. The following formula was used to calculate the irradiance:

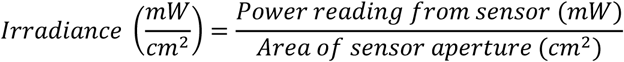

The temperature measurements at the C-chip were performed using a PT-100 (RTD thin film) element (TME® Electronic Components, Poland) connected to an electronic display. The PT-100 element was inserted between the layer of PDMS and glass forming the C-chip (Figure S3.c.i-ii.).

### 2.4. Cell culture

A cell line derived from human breast cancer, MDA-MB-231 (kindly provided by Dr. Marlena Godlewska; purchased from American Type Culture Collection), and mouse fibroblasts, L929, were used for biological studies. The cells were cultured in DMEM supplemented with 10% fetal bovine serum and a 1% mixture of antibiotics, penicillin-streptomycin, at 5% CO_2_ and 37°C in a humidified chamber. Cells were grown to confluence and harvested by trypsinization using a 0.25 mg/mL trypsin/EDTA solution and resuspended in fresh culture medium. Following Trypan Blue staining, viable cells were counted using the Countess II FL Automated Cell Counter. All cell culture reagents and materials were purchased from ThermoFisher (Waltham, MA, USA).

### 2.5. Hydrogel synthesis and sample preparation

Gelatin methacryloyl (GelMA) was synthesized according to the method described by Celikkin et al., 2022, with a few modifications. Shortly, 10 g of gelatin (Type A, Bloom 300; Sigma-Aldrich, United States) was completely dissolved in 400 mL of phosphate-buffered saline (Sigma-Aldrich, United States) at 50°C with continuous stirring. Methacrylic anhydride was added dropwise at a concentration of 0.8 mL per 1 g of dissolved gelatin to obtain a high degree of methacryloyl functionalization. The reaction was stopped after 5-6 h. The final solution was diluted with two volumes of preheated (at 40°C) PBS and transferred to a dialysis membrane (12-kDa MWCO, Sigma-Aldrich, United States) for 3-5 days. The solution was frozen at -80°C and transferred to the freeze dryer for 5 days. The freeze-dried solution was stored at -20°C.

HAMA (TintBionic® HAMA) and dECM from porcine pancreas (PrinTiss® dECM-PAN) were purchased from Polbionica sp. z o.o., Poland. The freeze-dried GelMA and/or HAMA powders were dissolved in PBS at the desired concentration by stirring for 1 h at 50°C. The dECM was added proportionally (0.5%) from a 5% dECM stock. The lithium phenyl-2,4,6-trimethylbenzoylphosphinate (LAP) photoinitiator was added at 0.4 % concentration by volume to the hydrogel solutions, and all prepared hydrogels were filtered. For cell encapsulation experiments, the cells were trypsinized from culture dishes, centrifuged, and resuspended in a fresh medium. Then, the cells were mixed with the hydrogel pre-polymer solution in 1:4 v/v (2.0 × 10^7^ cells/ml). The final hydrogel compositions used are as follows: (i) 4.8% GelMA, (ii) 4.8% GelMA + 0.4% TintBionic® HAMA (GelMA-HAMA), and (iii) 4.8% GelMA + 0.4% TintBionic® HAMA + 0.4% PrinTiss® dECM-PAN (GelMA-HAMA-ECM).

### 2.6. Hydrogel characterization

Droplets were generated using green fluorescent FITC-conjugated GelMA with the “on-the-fly” crosslinking system at 174, 884, and 1537 mW/cm^2^ irradiances. The droplets were imaged using a water-immersion 60x lens on the Nikon A1 confocal microscope. For imaging, the droplets were washed and fixed as a monolayer using low-melting-point agarose. The images were processed on ImageJ after generating a point spread function and deconvolution, using the BoneJ plugin for the pore size analysis.

Rheology analysis was done for 1 mL GelMA crosslinked into discs at the above mentioned light irradiances using the Anton Paar rheometer and the Anton Paar RheoCompass™ software. G’ (stress modulus) and G” (loss modulus) were measured after initial amplitude analysis to determine the applied strain range.

### 2.7. LIVE/DEAD cell staining and oxidative stress analysis

According to the manufacturer’s instructions, cell viability analysis was performed using a LIVE/DEAD kit. Briefly, encapsulated cells were seeded in multi-well plates and cultured for five days. On days 1, 3, and 5, the beads were collected in Eppendorf tubes, washed once with PBS, mixed with 30 μl of LIVE/DEAD kit stock solution (containing BOBO-1 Iodide and Calcein AM dyes) in DPBS (3:10), and incubated for 15 min at 37°C. Green fluorescence distinguished live cells based on enzymatic conversion of the non-fluorescent calcein AM to the fluorescent calcein inside the cells, while the red fluorescent dye penetrated only the dead/stressed cells with damaged membranes, binding to DNA. The beads were transferred to a microscope glass slide and observed using a green fluorescence channel (488 nm) for live cells and a red fluorescence channel (570 nm) for dead cells using Nikon Eclipse TSR2 (Nikon, Tokyo, Japan). Images were analyzed and visualized using the NIS-Elements software.

Oxidative stress in the encapsulated cells was analysed using CellROX™ Green Reagent. More precisely, 5 µM of the reagent was added to the droplet sample and incubated at 37°C for 30 minutes. The nuclei of the cells were stained with Hoechst 33342. The samples were observed under the microscope using the 405 nm (blue) and 488 nm (green) channels at 20× magnification. Green fluorescence was observed in ROS-affected cells. All kits were purchased from Thermofisher (Invitrogen, Waltham, USA).

### 2.8. Proliferation

To confirm the degree of proliferation of 3D-encapsulated MDA-MB-231 cells, we used the Cell Counting Kit-8 (CCK-8; WST-8[2-(2-methoxy-4-nitrophenyl)-3-(4-nitrophenyl)-5-(2,4-disulfophenyl)-2H-tetrazolium, monosodium salt; MedChem Express, United States). After cell encapsulation (using 1 mL of each hydrogel with 10 or 20 x 10^6^ cells), the beads were washed as described before and resuspended in 1800 µl of complete media. Next, 90 µl/well were seeded into a 96-well plate (in 5 replicates), and the rest of the cells were used for viability monitoring. The cells in beads were cultured for five days, and proliferation was checked after 1, 3, and 5 days of culture using 10 µl of CKK-8 reagent in each well and incubated for 4 h at 37°C. WST-8 was reduced by cellular dehydrogenases to form a water-soluble, orange-colored formazan product. The absorbance was measured at 450 nm using a Synergy HTX Microplate Spectrophotometer (BioTek, Winooski, VT, USA), and served as a measure of the number of living cells in the sample. For the long-term culture experiment, encapsulated cells were observed using a phase-contrast microscope for a period of 14 days (observed on Day 1, 3, 5, 7, and 14).

### 2.9. Statistical Analysis

We used GraphPad Prism 6.00 for Windows (GraphPad Software, San Diego, CA, USA) for statistical analysis of data. Data were analyzed using a non-parametric Mann–Whitney test. The *p*-value < 0.05 was considered statistically significant, with * *p* < 0.05, ** *p* < 0.01, and *** *p* < 0.001. For generating histograms, MATLAB Version: 9.13.0 (R2022b) Update 2 was used.

## 3. Results

### 3.1. Optimization of monodisperse droplet generation: flow rate and cell loading concentration

Microdroplets were generated using the D-chip by step-emulsification with different applied flow rates, *Q_H_* and *Q_C_* of the hydrogel phase (GelMA 4.8 % with 1x10^7^ MDA-MB-231 cells/ml or without cells) and the continuous phase (Novec®7500 with 2% PFPE-PEG-PFPE surfactant), respectively. We found that the presence of cells had little to no observable impact on the size *D* of the generated droplets (Figure 2.a.i-iii) for all tested rates of flow. We tested a fixed regime of hydrogel flow rates (*Q_H_* ∈ [6, 75] µL/min) at three different continuous phase flow rates (*Q_C_* = 25, 75, 125 µL/min).

**Figure 2.**
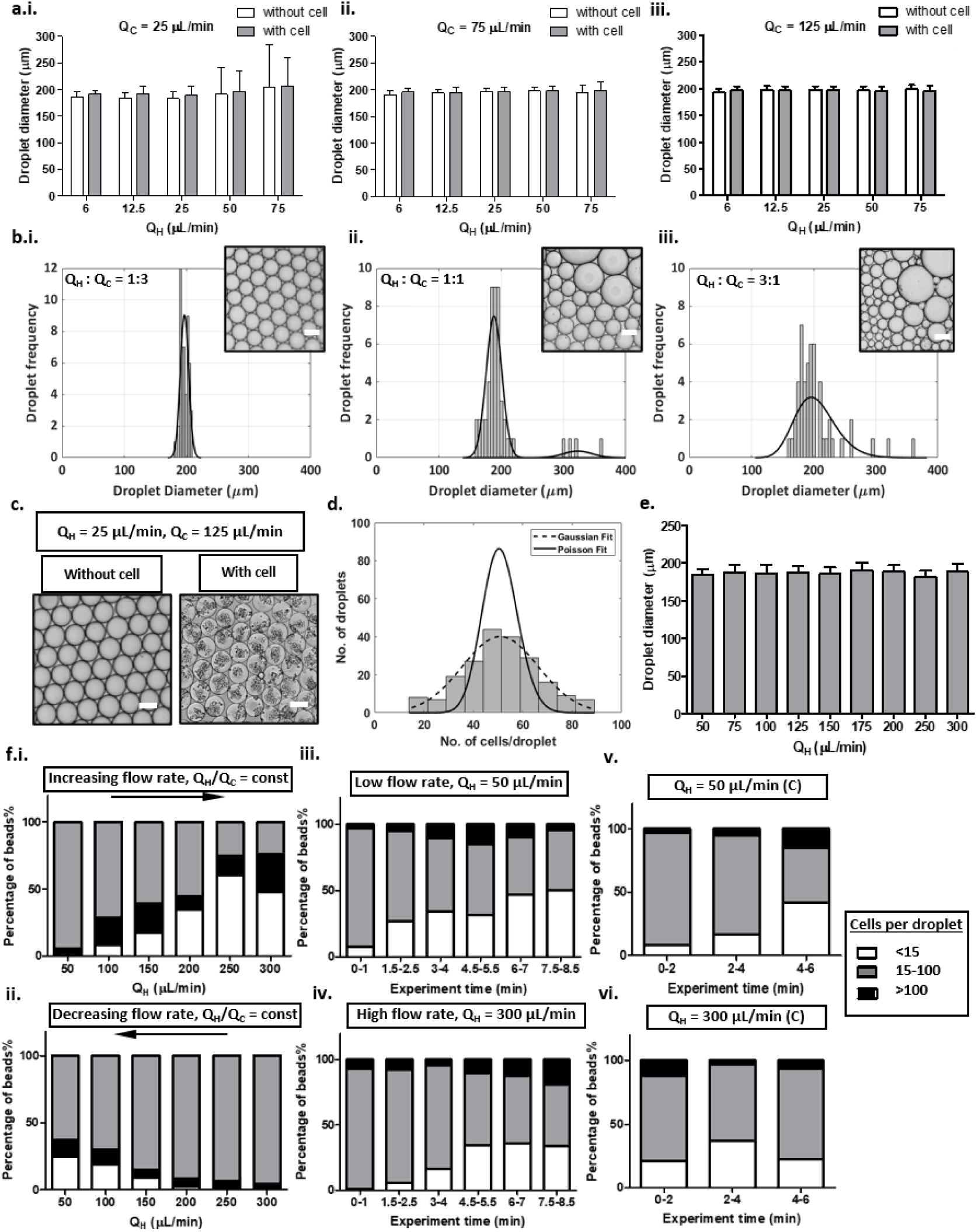
High-throughput droplet generation parameters. **a**. Average droplet diameters for hydrogel flow rates (Q_H_) (6, 12.5, 25, 50, and 75 µL/min) varied for continuous phase flow rates (Q_C_) of (i) 25, (ii) 75, and (iii) 125 µL/min. **b**. Histograms showing droplet diameter variations for different Q_H_ : Q_C_ volume fractions with Q_H_ - Q_C_ (i) 25-75, (ii) 50-50, and (iii) 75-25 µL/min for each case fitted to a Gaussian distribution curve (black line). **c.** Highly monodisperse droplets produced at Q_H_ : Q_C_ of 1:5 produced with and without encapsulated cells. **d.** Distribution curve for total cells encapsulated per droplet for a loading concentration of 20 million cells/mL fitted to Poisson (solid line) and Gaussian (dotted line) distribution curves. **e**. Average droplet diameter at high production rates at Q_H_ up to 300 µL/min. **f.** Changes in the encapsulated cell distribution: at Q_C_ /Q_H_=2=constant, for (i) increasing Q_H_, (ii) decreasing Q_H_; over time corresponding to 1-min experimental runs separated by 30s no-flow intervals with fixed Q_H_ of (iii) 50 µL/min and (iv) 300 µL/min; for droplets collected every 2 minutes for a continuous 6 min experimental run at Q_H_ – Q_C_ of (v) 50-100 µL/min and (vi) 300-600 µL/min.

We observed that the size and monodispersity of the droplets were nearly independent of *Q_C_* as long as *Q_C_* ≥ *Q_H_*. In this regime, the average droplet size *D* ranged from 185-200 µm, with slightly larger values observed at higher *Q_H_*/*Q_C_*, and with CV 3-4%, with the lowest CV values at the lowest studied *Q_H_*/*Q_C_*. In the cases with *Q_C_* < *Q_H_*, we observed a visible increase in the average size *D* with an increase in the coefficient of variation (CV) up to the level CV ∼ 30-40 % at the highest studied *Q_H_*/*Q_C_* (Figure 2.a. and Supplementary Table 1). In particular, we found that the distributions changed from strictly monodisperse to bidisperse and subsequently polydisperse as the ratio *Q_H_*:*Q_C_* increased from 1:3, via 1:1 to 3:1 at a fixed net flow rate *Q_H_* + *Q_C_* = 100 µL/min (Figure 2.b.i-iii). We attribute this widening of the distribution to droplet crowding in the collection channel near the outlet region. The presence of crowded droplets interfered with droplet generation and led, e.g., to delayed pinch-off or coalescence during or immediately post-generation.

In the cases with *Q_C_* ≥ *Q_H_*, we found that the most monodisperse droplets were typically observed at *Q_H_* = 25 µL/min, *Q_C_* = 125 µL/min, for which we found *D* = 198 μm, CV = 3.2% (Figure 2.c.). Very similar results, i.e., similar *D* and CV values, were also observed in droplets with encapsulated cells (Supplementary Table 1). In particular, at the same optimal set of flow rates (*Q_H_* = 25 µL/min, *Q_C_* = 125 µL/min), we found *D* = 198 μm, CV = 2.9% (Figure 2.c. and Supplementary Table 1.). From these observations, we conclude that the presence of cells had no visible impact on droplet size or monodispersity (at least at the applied cell concentrations in the range 10-20 x 10^6^/mL).

The above mentioned set of flow rates was also used to perform “on-the-fly” UV crosslinking, as described in the upcoming sections. The cell distribution per droplet was found to be Gaussian rather than Poissonian, which we attribute to moderate cell aggregation, with an average cell number per droplet, *N*= 50 for the cell concentration 20 x 10^6^/mL (Figure 2.d.). At a lower cell concentration of 10 x 10^6^/mL we observed more Poisson-like behaviour (Figure.S2.c), which may be related to limited aggregation in this case. The monodispersity in cell distribution was found to improve with increasing cell loading concentration, in agreement with the estimate for the Poisson distribution for which CV ∼ N^-1/2^.

Finally, in a quest for the highest achievable throughput, we verified operation of the D-chip at even higher *Q_H_* values ranging from 50 to 300 µL/min (Figure 2.e.), while keeping *Q_C_* = 2*Q_H_*, i.e., such that the continuous phase flow rate *Q_C_* was always large enough to warrant droplet monodispersity but also not excessive to limit oil consumption. We found that the size of the droplets remained roughly constant and monodispersity could be safely retained at the level CV = 5-6 % for hydrogel flow rates up to *Q_H_* = 300 µL/min, which corresponds to the production rate of up to ∼95 000 droplets/min (Supplementary Movie 1). We note that perhaps even higher rates of flow could be achieved since we did not observe any issues related to droplet monodispersity. The only possible limitation for the droplet generation rates could be the mechanical stability of the chip itself, since with increasing flow rates the pressure inside the chip also increased. In cases with poor bonding of PDMS channels to the glass slide, we observed PDMS delamination and leakage. However, with good bonding, the D-chip was capable of achieving even higher production rates.

### 3.2. The effect of flow rate and experimental protocol on cell distribution

During experiments with cells, we observed that cell distribution in droplets seems to be less uniform at lower Q_H_ and that, in cases with continuous droplet generation, it further seems to deteriorate during the experiment on the scale of minutes. In order to investigate this phenomenon in more detail, we decided to systematically study the impact of the applied rates of flow and their changes in time on the observed cell distribution (Figure 2.f.). First, we tested the impact of *Q_H_* while setting *Q_C_* = 2*Q_H_*. The experimental protocol consisted of 6 experimental runs with *Q_H_* gradually increasing from 50 to 300 µL/min (Figure 2.f.i). Each run started with 1 min of constant flow rate at fixed *Q_H_* followed by a 30s no-flow period. During the first run at *Q_H_* = 50 µL/min, we found the distribution to be uniform, with >90% of droplets having the occupancy close to the expected mean value, but then the distribution deteriorated already in the second run (*Q_H_* = 100 µL/min) and further deteriorated at subsequent runs. In the last run we observed less than 25% of droplets having the expected cell occupancy, with remaining droplets either almost empty or consisting of strongly aggregated cell clumps. However, it was not clear whether the dominant factor responsible for the aggregation—and so deterioration in the distribution—was the flow rate *Q_H_* or the duration of the experiment. To check this, we inverted the order of the experimental runs, i.e., so that *Q_H_* decreased from 300 to 50 µL/min (Figure 2.f.ii). In this case, we observed uniform distribution for the first 3 runs, and while gradual deterioration followed from run 4, even in run 6 we found that > 50% of droplets had the expected occupancy. Accordingly, we have established that the higher flow rates tend to delay cell aggregation. These results overall suggest that (i) the deterioration of the cell distribution is a time-rather than flow-rate dependent process, and that (ii) higher flow rates seem to reduce this unwanted effect (Figure S2.b.).

To further decouple the impact of flow and time, we performed a second series of experiments in which we applied a similar protocol consisting also of 6 runs (each run being 1 min flow + 30 seconds no-flow), but with all runs performed at the same fixed *Q_H_* (Figure 2.f.iii. and iv.). We tested *Q_H_* = 50 µL/min and *Q_H_* = 300 µL/min, and while in the latter case the uniformity was retained for a longer time (up to the third run), in both cases we found gradual loss of uniformity. Finally, we also performed a series of 3 runs of a constant flow at a fixed *Q_H_*, with each run lasting 2 minutes, i.e., *without* the no-flow intervals (Figure 2.f.v. and vi.). While at *Q_H_* = 50 µL/min (Figure 2.f.v.) we found a result similar to the previous cases (<50% expected occupancy in the last run), at *Q_H_* = 300 µL/min the uniformity of the distribution was roughly retained throughout all runs (Figure 2.f.vi). In particular, the expected occupancy stabilized at around 70% in the last run. The fact that uniformity in the first run was lower than in previous experiments may be attributed to pre-aggregation of cells inside the inlet tubing, which in this case (i.e., in continuous 6 min flow at *Q_H_* = 300 µL/min) was much longer (around 100 cm long) than in other experiments. Overall, we conclude that the highest studied flow rates (and only those) warrant uniform cell distributions at long device operation times. This could be seen as particularly advantageous for high-throughput applications requiring long-term continuous operation.

Finally, we attempt to rationalize the ‘uniformization’ of the distributions of the high applied *Q_H_*. We propose that this effect relies on flow-mediated prevention of sedimentation and aggregation of cells on the D-chip. Indeed, a sufficiently fast-flowing hydrogel can be expected to (i) induce cell mixing due to hydrodynamic lift forces which exceed gravitational forces^53^ and (ii) impose shear stresses which disperse pre-formed cell aggregates into single dissociated cells. To further improve the uniformity, one could use flow-based homogenizers, e.g., flow chambers consisting of narrowly spaced pillars, wherein viscous stresses dissociate the aggregates^54^.

### 3.3. Variations of droplet size distributions in different droplet generators

The 64 parallel droplet generators were numbered as indicated in Figure 3.a. The angular positions of each of the channels were determined based on their position with respect to the inlet axis (connecting the oil and hydrogel inlets). The droplet generation frequency from each of the 64 droplet generator channels was measured for Q_H_ = 25 µL/min and Q_H_ = 50 µL/min (Figure 3.b.). In general, we find quite significant variations in the frequency of droplet generation between individual channels, with CV_freq_ of 12.5% (at Q_H_ = 50 µL/min) and 17.8% (at Q_H_ = 25 µL/min). These fluctuations are even more pronounced in the process involving cells, with CV_freq_ = 27.5% (at Q_H_ = 50 µL/min) and 43.8% (at Q_H_ = 25 µL/min). The CV decreases as the flow rates are increased. This can be attributed to cell aggregation in some channels, resulting in higher hydrodynamic resistance and lower local flow rates, as well as a positive feedback loop for cell aggregation (local lowering of the flow rate enhances aggregations). In all cases, decreased generation frequency could be observed at the generators closest to the outlet, which can be attributed to the droplet crowding effect in this region.

**Figure 3.**
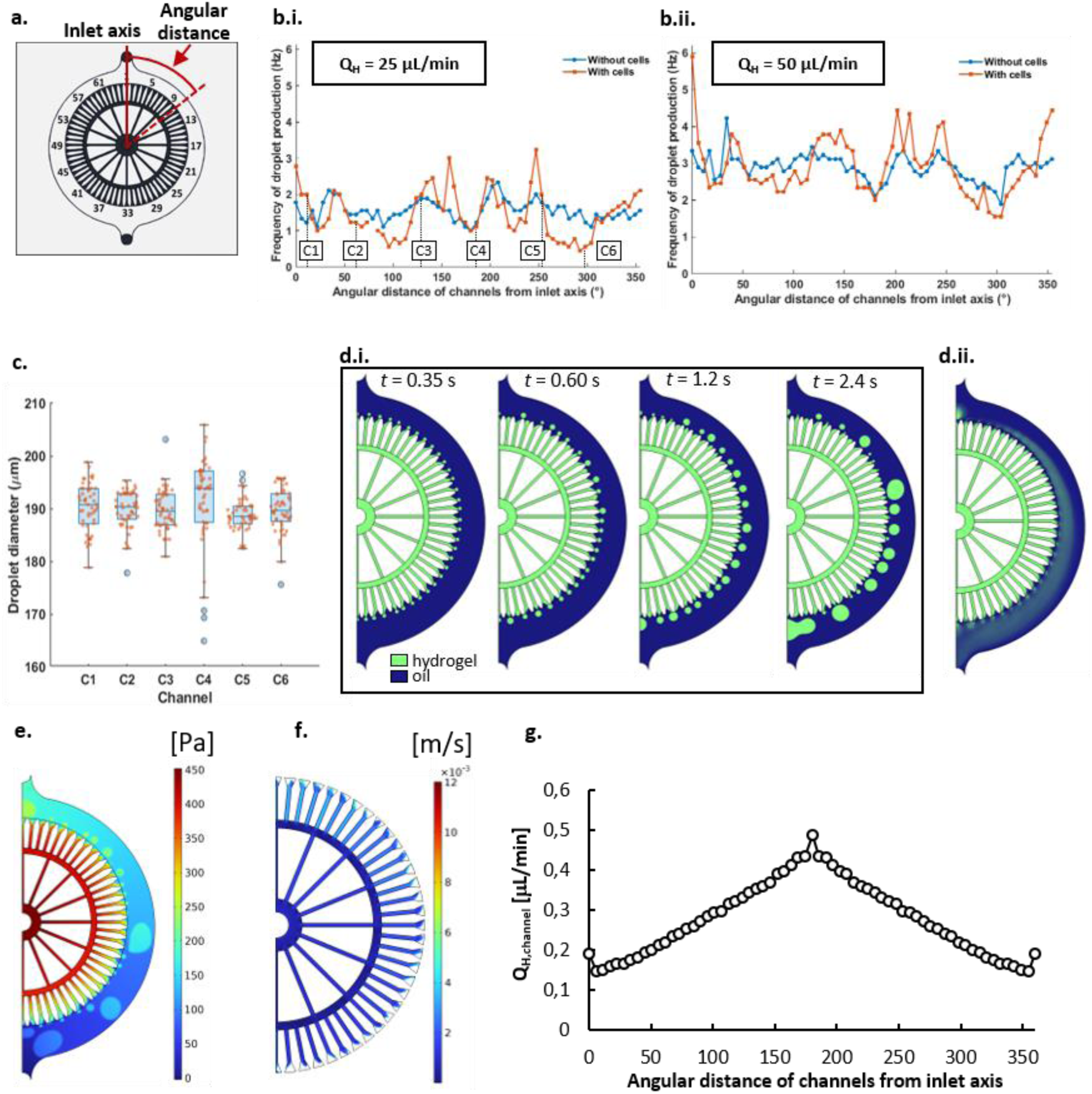
Droplet production in different microchannels. **a.** Layout showing the numbering of droplet generation channels and defining their angular positions; **b.** Frequency of droplet generation measured in each of the 64 channels at Q_C_ = 125 µL/min, with and without cells for i. Q_H_ = 25 µL/min with 6 selected channels marked C1, C2, C3, C4, C5, and C6 representing channels 3, 12, 24, 34, 46, and 54, respectively; and ii. Q_H_ = 50 µL/min; **c.** Size variations observed among droplets (with cells) produced from the 6 selected channels at Q_H_ = 25 µL/min and Q_C_ = 125 µL/min. **Numerical simulation studies**. **d.i.** Snapshots depicting the process of droplet formation and coalescence within the first 2 seconds after the onset of the flow (hydrogel and oil phases indicated in green and blue colors, respectively). **d.ii.** Distribution of the time-averaged hydrogel volume fraction inside the device. **e.** Pressure distribution in the chip at t = 20 s. **f.** Velocity magnitude in the microchannels at t = 20 s. **g.** Mean volumetric flow rate Q_H,channel_ of the hydrogel inside individual microchannels as a function of the angular distance from the inlet axis.

The height of the step, *H*_S_, and the height of the droplet generation channel, *H*_C_, were measured for each generator using optical profilometry. The average measured values were *H*_C_ = 43 ± 7 µm and H_S_ = 182 ± 12 µm (Figure S1.f). The variations in the measured *H*_C_ and *H*_S_ at individual droplet generators were mostly due to micromilling error margins during the chip fabrication. Next, 6 droplet generator channels were selected (C1-channel 3, C2-channel 12, C3-channel 24, C4-channel 34, C5-channel 46, and C6-channel 54) from different regions of the chip. Droplet generation from each of these channels was monitored, and droplet diameters of 60 droplets generated from each channel were measured (corresponding droplet size distributions are shown in Figure S2.d). We found that the channel with the maximum CV in droplet sizes was channel C4, which is closest to the outlet. Again, the large CV in this case could be attributed to droplet crowding in this region.

### 3.4. Numerical simulations of droplet generation

In order to better understand the distribution of the microflows and the dynamics of droplet formation inside the droplet-generating chip (D-chip), we pursued 3D numerical simulations of the two-phase fluid flow using the finite element method, which we have implemented via COMSOL software. Our simulations covered 20 seconds of the operation of the device and resolved the interface between the two immiscible phases forming inside the device, as well as the Young’s contact angle *θ*_Y_ of the droplet phase at the walls of the channels, which we set *θ*_Y_ = 179° to impose the non-wetting conditions. The simulations did not incorporate any type of surfactant in the oil phase, so the simulated droplets coalesced quickly after being formed. However, we could still extract useful information about the operation of the device, such as the pressure distribution and the velocity field inside the microchannels.

The simulation starts with a configuration in which the hydrogel phase occupies the inlet, the distribution channels, and the droplet-generating microchannels, whereas the oil phase fills the circular droplet collection channel. The two phases are separated by an interface forming along the inner rim of the collection channel. Snapshots in Figure 3.d.i depict the first 2 seconds of the operation of the device following the onset of the flow. First, we observe the formation of a droplet at each of the junctions. Those droplets feature a rather monodisperse size distribution. However, in the following snapshots, we observe droplet coalescence resulting in polydispersity. In addition, in some cases, the droplets forming at a given microchannel coalesce with the advancing hydrogel-oil interface inside the same microchannel, which effectively disables droplet pinch-off and results in the formation of a growing hydrogel ‘bulge’ (eventually advected by the flow as a huge droplet). Therefore, direct counting of the forming droplets at individual generators appears unfeasible. Instead, we extract the volume fraction of the hydrogel phase averaged out over the duration of the whole simulation (Figure 3.d.ii), as well as the instantaneous distribution of the pressure (Figure 3.e.) and the flow velocity (more precisely, the absolute value of the velocity vector field; Figure 3.f.) inside the device at t = 20 s. As expected, we find that the pressure inside the circular collection channel gradually decreases towards the outlet (Figure 3.e.). This leads to an increasing pressure difference between the inlets and the outlets of the individual droplet generators, and consequently to a gradually increasing flow velocity inside microchannels while moving downstream in the collection channel (Figure 3.f). As a result, we find a roughly linearly increasing time-averaged rate of flow *Q_H_* inside the microchannels as a function of their angular position along the perimeter of the circular collection channel measured towards the outlet (Figure 3.g). The data in the angle range 180°-360° mirror the data from the range 0°-180° as we assume periodic boundary conditions along the inlet axis (reflecting the symmetry of the device).

From the known mechanism of droplet formation at a step-junction as demonstrated, e.g., by Eggersdorfer *et al.*, we may expect the droplet size to be nearly independent of the applied rates of flow^55^. Therefore, based on the extracted average flow rates *Q_H_* inside the microchannels (Figure 3.g), one would expect the droplet frequency to proportionally *increase* towards the outlet - by a factor of around 2.5. This expectation, however, remains in stark contrast with the droplet generation frequencies measured in the experiment (Figure 3.b.) wherein we do not observe any such trend, but instead, as already mentioned, we find a local minimum in the droplet generation frequency close to the outlet (180°). A possible explanation is that, in the experiment, droplet crowding (which we empirically observe) leads to their direct-contact interactions, such that the newly forming hydrogel-oil interfaces at the microchannel outlets experience a back-pressure (additional Laplace pressure) in this region. This could result in a locally lowered hydrogel flow rates and decreased droplet production rates, as observed in the experiment. However, in order to confirm this hypothesis via simulations, one would need to resort to a more advanced model incorporating the presence of surfactant (or providing another mechanism preventing droplet coalescence). We leave such studies as possible future work.

### 3.4. Effect of photocrosslinking efficiency on droplet structure

Photo-crosslinking of the microdroplets was evaluated using three lamps: (i) the home-made lamp (‘HL’) with LEDs of wavelength 405 nm (94-255 mW/cm^2^), (ii) the commercial lamp (‘CL’) of wavelength 385 nm (174-1537 mW/cm^2^) and (iii) the commercial lamp (‘CL’) with wavelength 405 nm (174-1537 mW/cm^2^). The lamps were characterized in terms of irradiance and light-induced heat generated on-chip (we measured the temperature on-chip); see Figure S3.a-d. For droplet crosslinking, the lamps were placed at a distance of 0.5 cm from the C-chip to obtain a possibly uniform, high irradiance at a given input current (Figure S3.b).

We found that the irradiances in the range of 90-250 mW/cm^2^ were insufficient for complete crosslinking of the GelMA droplets (4.8% w/w GelMA in PBS). The incomplete crosslinking was signified by the formation of a visible core-shell structure (Figure S4.a. and b. and Figure 4a.ii). This observation was in line with the previous report by the Oakey group^56^, who attributed the emergence of the core-shell structure to poor crosslinking of the region close to the hydrogel-oil interface associated with the diffusion of photopolymerization-inhibiting oxygen dissolved in the oil phase. Indeed, fluorinated oils such as Novec 7500 typically have high oxygen solubility, while PDMS is also gas-permeable. We observed the emergence of the core-shell structure in both cases with 405 nm and 385 nm illumination wavelengths. However, the shell thickness *t* was found to be consistently smaller in the latter case (except at the maximum irradiance around 1500 mW/cm^2^, for which *t* was similar in both cases); see Figure 4.a.i-ii. At intermediate irradiances, in the range 250-1300 mW/ cm^2^, we observed that, in the case of 385 nm light, the shell thickness developed a well-defined minimum at around 884 mW/cm^2^. At such optimal irradiance, the shell thickness went down to around or below 15 microns, i.e., close to the detection limit via conventional transmitted light microscopy (which we used in this case), and in many cases the shell was not visible at all, suggesting complete (homogeneous) crosslinking. In the case of the 405 nm light, the shell thickness remained approximately constant, independent of the irradiance and equal to ∼30 µm, i.e., the beads were never completely crosslinked in this case. The presence of “shells” in the incompletely crosslinked droplets could be seen both with and without cell encapsulation (Figure 4.a. and S4.b.).

**Figure 4.**
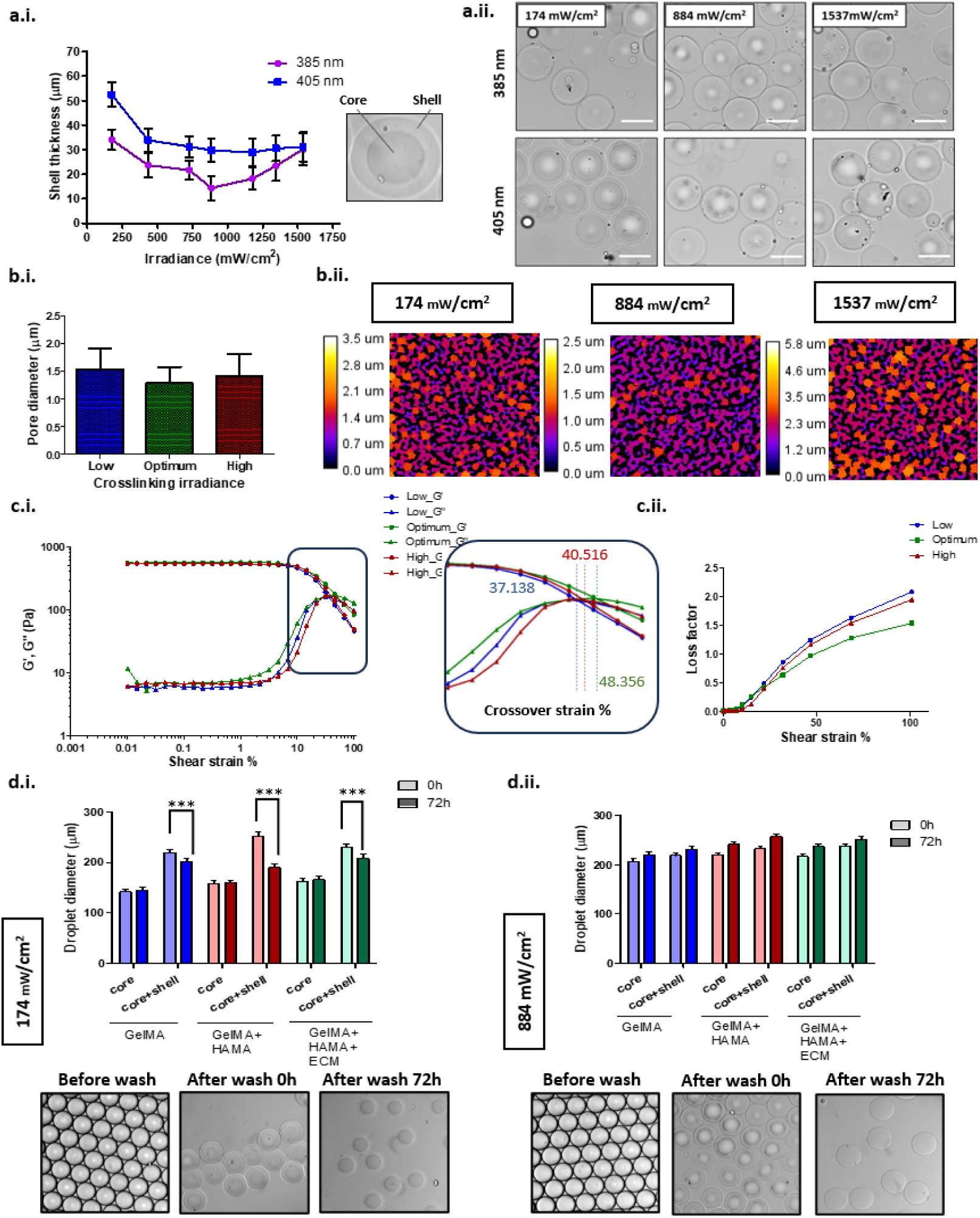
Effects of photo-crosslinking on droplet structure. **a.i.** A graph showing variations in the shell thickness based on UV irradiance and wavelength used for crosslinking with inset showing an incompletely crosslinked droplet with a core-shell structure. **ii.** Variations in droplet structure seen at low, optimum and high irradiances (174, 884, and 1537 mW/cm² respectively) for the two different wavelength CLs. **b.** BoneJ analysis of the pore size distribution of FITC-GelMA droplets crosslinked at low, optimum and high irradiance with **i.** a graph showing the average pore diameters of the crosslinked structures and **ii.** fire maps for pore size distributions for sections from the droplets analyzed using confocal microscopy. **c.** Rheological analysis of bulk GelMA hydrogels crosslinked at the low, optimum, and high UV irradiances showing **i.** Storage (G’) and loss (G”) modulus variations with inset showing the crossover strain percentages in each case, and **ii.** loss factor variations, for bulk hydrogels (1 mL/sample). **d.** Presence of shell seen in GELMA + HAMA droplets seen after crosslinking at **i**. low (174 mW/cm²) vs **ii.** optimum irradiance (884 mW/cm²) showing droplet size reduction seen over time (72h) in the former case due to dissolution of the shell while droplet size increased for well crosslinked droplets at optimum irradiance (Scale = 200 μm; P-value: *-0.05, **-0.01,***-0.001).

Furthermore, the droplets crosslinked at irradiances above 1000 mW/cm^2^ were found to be less stable (up to about 72 h) than those crosslinked at irradiances below 1000 mW/cm^2^. Temperature at the irradiation spot rose even above 60°C at these irradiance levels (Figure S3.c.ii). Even though this exposure was not enough to heat up the chip or the continuous phase (Figure S3.c.i and d), it could affect the reaction kinetics and physicochemical properties of the hydrogel, thereby affecting its stability^57^.

To study the effect of irradiance on the degree of crosslinking, we additionally generated FITC-GelMA droplets using the minimum (174 mW/cm^2^), optimum (884 mW/cm^2^), and maximum (1537 mW/cm^2^) irradiances (termed as “low”, “optimum”, and “high”, respectively). BoneJ analysis of the droplets revealed the “optimum” sample to have the lowest average pore size and minimum heterogeneity (1.3±0.3 µm) among the three samples (Figure 4.b.i). The “low” irradiance sample had the highest average pore diameter (1.5 µm), while the “high” irradiance sample had the most heterogeneous pores, with the largest pore reaching around 5.8 µm (Figure 4.b.ii). The hydrogel pore features can be attributed to the effect of irradiance level on the photopolymerization reaction kinetics. In the case of low irradiance, lesser number of photoinitiator molecules are cleaved, resulting in a lower number of polymer chains to be activated and, as a result, longer polymer chains to be formed prior to termination^58^. On the other hand, at higher irradiances, several photoinitiator molecules are cleaved, leading to a rapid crosslinking reaction, happening faster than the diffusion of the unreacted monomers, resulting in short, dense chains forming at particular regions with interspersed loose, irregular voids surrounding them^59^.

We complemented the microporosity measurements with rheological analysis of the viscoelastic properties of the bulk hydrogel discs at similar crosslinking conditions (yet off-chip and without the presence of the oil phase). The analysis revealed a more flexible architecture at the “optimum” irradiance, evidenced by the highest crossover strain, meaning the highest strain tolerance compared to the “low” and “high” samples^58,60^ (Figure 4.c.i.). Furthermore, the “optimum” sample also had the lowest loss factor (tan δ= G’’ / G’), signifying a more elastic solid with lower damping capacity^61^ (Figure 4.c.ii.). The optimally crosslinked droplets were then found to be stable for at least two weeks in cell culture conditions (37°C, 5% CO_2_, 85% humidity).

To check if the crosslinking conditions changed depending on the hydrogel composition, droplets of GelMA, GelMA-HAMA, and GelMA-HAMA-ECM were crosslinked at “low” and “optimum” irradiance (Figure 4.d. and S4.c.). In previous studies, HAMA has been shown to improve the structural durability of the hydrogel and long-term viability of the encapsulated cells, while the ECM (dECM) has been shown to provide essential growth factors and improve cell growth as compared to bare non-protein-based or non-tissue-specific matrices (such as pure collagen)^31,62^. In all three cases, the presence of the under-crosslinked shell was observed for the “low” irradiance condition. The shell dissolved over a time period of 72 hours, resulting in a significant decrease in the bead diameter. On the other hand, the droplets crosslinked at “optimum” irradiance not only retained their size but also swelled and slightly increased in diameter within 72 hours. Therefore, our system could be used at the “optimum” crosslinking conditions with different hydrogel compositions. However, optimization of crosslinking conditions might be needed if the base hydrogel and/or crosslinking group or photoinitiator were changed.

### 3.5. Crosslinking efficiency affects cell viability

Next, we checked the viability of the encapsulated cells in the photocrosslinked droplets under 385 nm light exposure. Mouse fibroblasts (L929) were encapsulated in 4.8% GelMA + 0.4% HAMA droplets and exposed to different irradiances. The LIVE/DEAD staining of the encapsulated cells on Day 1 of culture revealed cell viability above 80% for all irradiances less than 1000 mW/cm^2^. The viability declined gradually for the droplets crosslinked at higher irradiances (Figure 5.a.i. and ii.).

**Figure 5.**
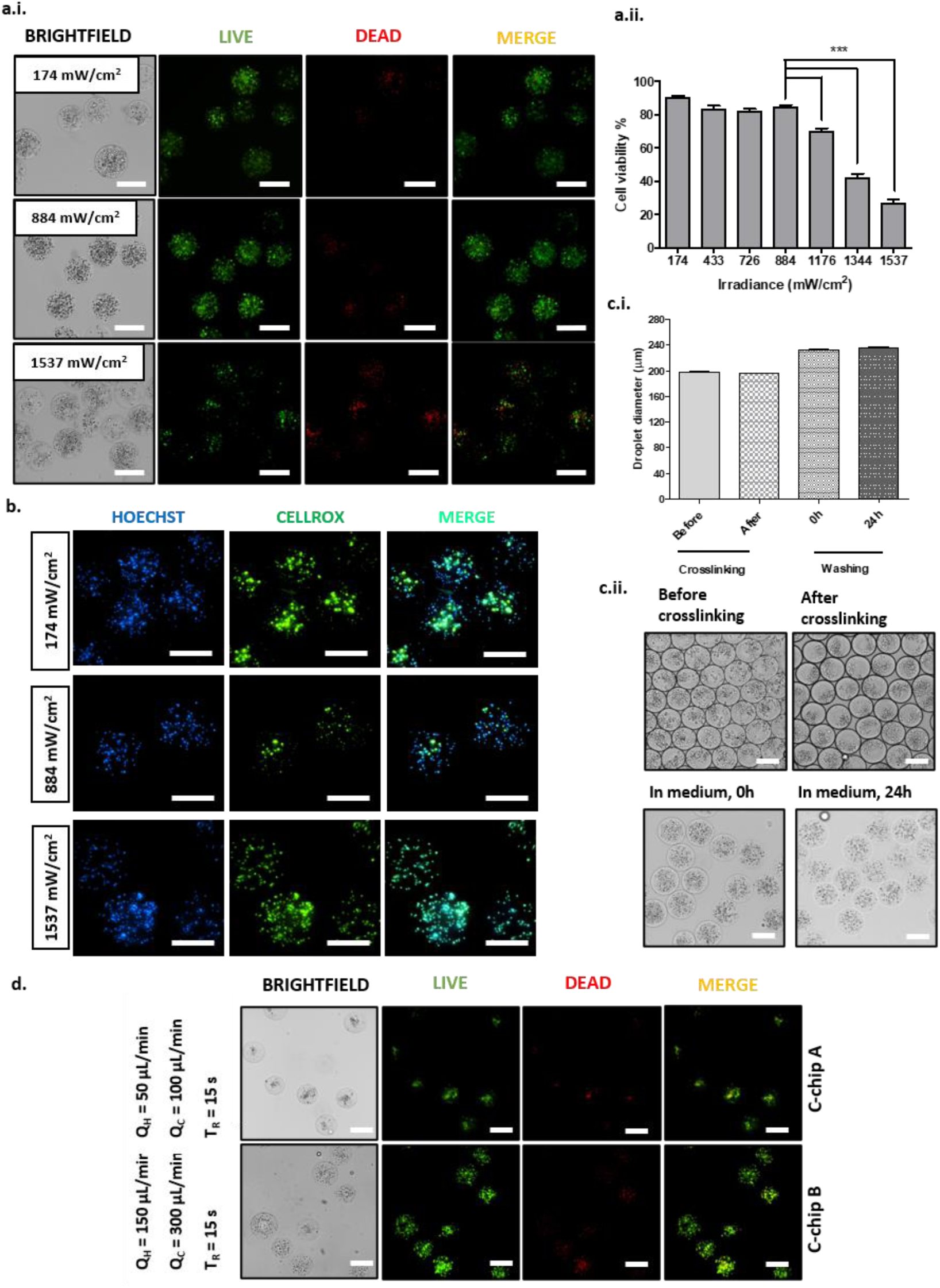
Analysis of cell viability in encapsulation [Day 1 of culture]. **a.** Viability of encapsulated L929 cells in 4.8% GelMA with 0.4% HAMA **i.** using live (green) – dead (red) fluorescent assay for low, optimum and high irradiance (174, 884 and 1537 mW/cm² respectively), and **ii.** Percentage of viable cells seen for droplets crosslinked using different irradiances of the 385 nm lamp. **b.** Analysis of oxidative stress using CellRox (green) assay (nucleus counterstained with Hoechst 33342 (blue) for droplets crosslinked at low, optimum, and high irradiance. **c.** Well crosslinked droplets seen at different stages of production with **i.** their diameter and **ii**. structural appearance during each stage. **d.** Cell viability observed by live-dead analysis at higher flow rates to increase droplet production rate by 2X (top) and 6X (bottom) crosslinked at equal residence time (T_R_) using C-chips with different channel lengths (Scale = 200 µm; P-value: *-0.05, **-0.01,***-0.001).

The samples crosslinked at low, optimal, and high irradiances (174, 884, and 1537 mW/cm^2^, respectively) were checked for the presence of reactive oxygen species (ROS) via the CellRox assay as indicative of oxidative cell stress, e.g., due to the UV crosslinking process. The results showed that the sample crosslinked at the optimal irradiance had the least number of cells showing oxidative stress as compared to the low- and high-irradiance samples (Figure 5.b.). This was further validated with a Live/Dead assay at Day 3 (Figure S5.a.ii.), which showed that viability further declined in the case of the low-irradiance sample. Additionally, the dissolution of the interfacial hydrogel layer, the ‘shell’, could also be observed in these droplets, as reported in the previous section. This, in turn, resulted in the release of the encapsulated cells. The possible reason for high levels of ROS (reactive oxygen species) could be due to the presence of active free radicals in the incompletely crosslinked “shell” and also due to the direct effect of high-irradiance UV light.

In summary, crosslinking with the 385 nm CL at an exposure time of 15 seconds was found to be the most efficient in terms of microbead stability and cell viability. The viability of cells was found to be around 85 % (±1%) on day 1 of culture and further increased on day 3. Around a 30% increase in the number of cells per droplet at day 3 compared to day 1 was observed under this condition, indicating active proliferation. We used the above-mentioned “optimal” crosslinking conditions (884 mW/cm^2^, 385 nm light) in all further experiments. The beads were carefully monitored at each step of the encapsulation process: (i) production, (ii) crosslinking, and (iii) washing (Day 0 and 1) (Figure 5.c.i). The size of the produced (non-crosslinked) microdroplets was measured to be *D* = 198 ± 6 µm, while the size of the crosslinked microbeads was found to be *D* = 195 µm ± 6 µm. Immediately after washing and transferring to cell culture medium, the droplet size increased to 232 ± 7 µm, which we attributed to hydrogel swelling due to contact with the external aqueous phase. We did not observe any significant further swelling after 24 h of culture in medium when the average droplet size was measured to be *D* = 235 ± 10 µm (Figure 5.c.ii).

To check whether even higher production rates could be achieved with the “on-the-fly” crosslinking system, the flow rates of *Q_H_*-*Q_C_* of (i) 50-100 µL/min and (ii) 150-300 µL/min were attempted with 4.8% GelMA. Cell viability in cases (i) and (ii) was found to be ∼68% and ∼71% at Day 1 of culture (Figure 5.d.). Since these values were slightly lower compared to the ∼76% obtained in case (iii) with *Q_H_*-*Q_C_* of 25-125 µL/min, we selected the latter set of flow rates to carry on the experiments with cells. In any case, as can be seen, the higher production rates were also achievable with our system at a reasonable cell viability. In particular, the maximal demonstrated crosslinked microbead production rate with encapsulated cells (achievable at Q_H_ = 150 uL/min) was ∼45 000 microbeads/min.

### 3.6. Hydrogel composition modulates cell growth

To check for cellular aggregation (an essential feature in tumor modelling), we used breast cancer cell line MDA-MB-231 with genetic characteristics representative of triple-negative breast cancer. We added the cells at a concentration of 20 million cells/mL using three different hydrogel compositions: GelMA, GelMA-HAMA, or GelMA-HAMA-ECM. Microdroplets were generated using our ‘on-the-fly’ crosslinking system at high throughput and cultured for 5 days to check for proliferation and microtissue formation (Figure 6.a. and S5.b.). In GelMA, around 75% of cells were alive after 24 h of culture in microcapsules; however, MDA-MB-231 cells, after five days, digested the matrix structure and migrated out of the microbeads (Figure S5.b). In GelMA-HAMA, the hydrogel containing ECM showed improved viability of the encapsulated cells, and the majority (around 75% in GelMA, 80 % in GelMA-HAMA, and 90 % in GelMA-HAMA-ECM) of tested cells were alive after one day of culture (Figure 6.a.ii). In the latter two cases, the microbeads were stable, and the cells were trapped inside the beads for the entire duration of the experiment. Presumably, the interpenetrating networks of the three types of matrices may have prevented rapid cell escape. In the examination of cell growth using CCK-8, we also established that the proliferation was significantly (\**p<* 0.05) higher in the GelMA-HAMA and GelMA-HAMA-ECM beads compared to pure GelMA beads(Figure 6.a.iii). Altogether, the GelMA-HAMA-ECM sample was found to show the highest cell viability and the fastest cell growth by Day 5 compared to the other two hydrogel compositions.

**Figure 6.**
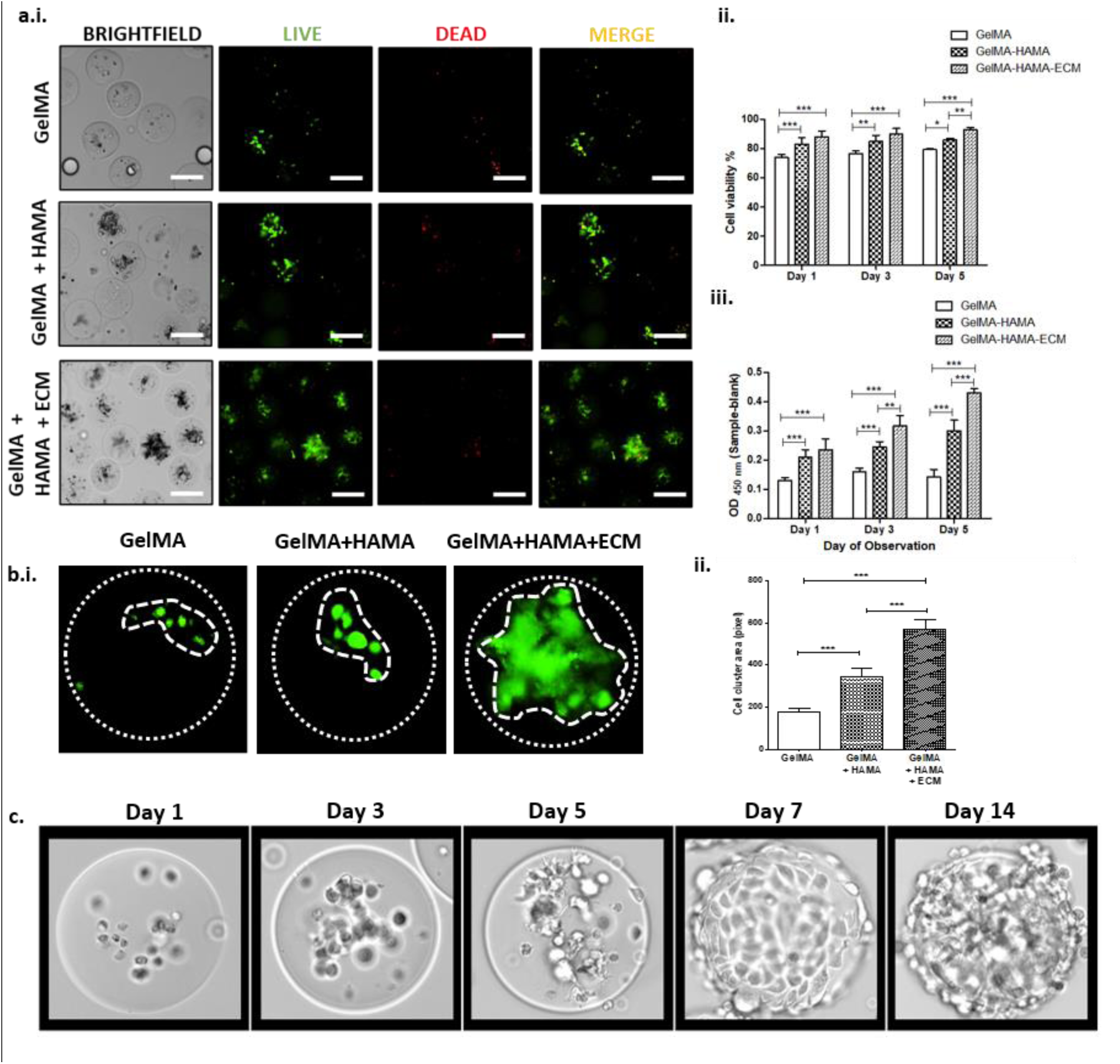
Cell growth depending on hydrogel composition. **a.** Viability of breast cancer cells MDA-MB-231 encapsulated in 4.8 % GelMA, 4.8 % GelMA+ 0.4 % HAMA, and GelMA 4.8 % with 0.4 % HAMA and 0.4 % ECM **i.** observed after LIVE/DEAD staining on Day 1 of culture (green and red fluorescence indicating live and dead cells respectively) and **ii.** percentage viability of the cells and **iii.** cell proliferation observed by CCK-8 assay till 5 days of culture after encapsulation. **b.** Cell clustering/spheroid formation **i.** structure and **ii.** sizes observed with Calcein-AM staining after 5 days of culture in different hydrogels. **c.** Long-term culture of L929 fibroblasts in GelMA-HAMA-ECM hydrogel upto 14 days showing microtissue formation. (Scale = 200 μm; P-value: *-0.05, **-0.01,***-0.001).

Furthermore, the GelMA-HAMA-ECM composition additionally stimulated the cells for enhanced aggregation (Figure 6.b.i). The aggregates created in the ECM-containing hydrogel were 2x-3x larger than in GelMA and GelMA-HAMA hydrogels and formed faster. By Day 5, cells in GelMA-HAMA also showed significant aggregation (Figure S5.b.).

To expand the portfolio of potential microtissue engineering applications, we have also checked the effect of the ECM-containing hydrogel on the growth of encapsulated L929 fibroblasts. Cells were encapsulated at a concentration of 10 million cells/mL in droplets containing a GelMA-HAMA-ECM hydrogel mixture. The microbeads were found to be stable in culture for up to 14 days (Figure 6.c.), and rapid proliferation of the cells was observed. The fibroblasts behaved differently from MDA-MB-231 cells and migrated onto the microbead surface, where they continued to grow, generating spherical microtissues after 14 days.

Overall, we may conclude that the addition of both HAMA and ECM components represents a potentially significant advantage in cell modelling or drug testing applications based on encapsulated microtissues.

## 4. Discussion

In this work, we demonstrate a microfluidic platform for high-throughput production of monodisperse, cell-laden GelMA microbeads suitable for scalable multicellular microtissue formation. Critically, the system overcomes the challenge of combining high cell loading with high-throughput encapsulation by generating ∼200 µm sized beads containing several tens of cells, followed by controlled photocrosslinking to ensure production of highly durable microbeads with good cell viability. The platform combines-i. the droplet generator, D-chip, that enables rapid dispersion of a hydrogel-cell mixture into monodisperse droplets at rates up to ∼95 000 droplets/min using 64 parallel droplet generators (step-emulsifiers) that warrant robust droplet production at relatively low viscous stresses, high droplet monodispersity, and good cell viability post-encapsulation; and ii. the photocrosslinking platform, C-chip, with a narrow, meandering microchannel (width only slightly exceeding the droplet diameter) to ensure no droplet-channel contact, allowing high fidelity of spherical droplet structure at high flow rates while allowing adequate residence times (*T_R_*) for UV-Vis light-driven photocrosslinking in flow (“on-the-fly” crosslinking). We fine-tune the flow rates, length of the channel, light wavelength, irradiance, as well as hydrogel composition to achieve optimal microbead stability, cell viability, and microtissue formation within up to 14 days in culture.

Optimization of the crosslinking conditions was crucial for maintaining both bead stability and cell viability. We observed a core-shell structure under some irradiance conditions, with partial shell dissolution after incubation in PBS/media, indicating incomplete crosslinking near the droplet interface, likely associated with oxygen inhibition of photopolymerization^56,63^. We further establish that:

i. For the case with the wavelength 385 nm, the shells are significantly thinner than at 405 nm, possibly due to better wavelength matching with the absorbance spectra of LAP photoinitiator peaking at around 380 nm^64^.
ii. In the case of 385 nm, the shell thickness *t* depends non-monotonically on the irradiance in the range 170-1600 mW/cm^2^, with a well-defined minimum *t*_min_∼12 µm (being close to the limit of optical detection in our case) at the optimized irradiance of around 900 mW/cm^2^.
iii. Increasing the irradiance beyond the optimum 900 mW/cm^2^ leads to significantly reduced cell viability while maximum structural stability (more homogeneous and smaller pores seen in BoneJ analysis) at the optimum irradiance is warranted by better viscoelastic properties (lower loss factor and higher crossover strain)^65^.

The existence of an optimal irradiance is particularly relevant for high-throughput photocrosslinking. Although increased irradiance generally accelerates photopolymerization^66^, the high intensities, required here for sufficiently fast crosslinking, generate substantial amount of heat, with local PDMS temperatures increasing by up to ∼60 °C above 1000 mW/cm², even under imposed air circulation (via electric fan). Increased temperature can enhance oxygen diffusion and thereby counteract polymerization, while also altering hydrogel network properties^67–69^. Although the precise mechanism underlying the observed non-monotonic behaviour requires further investigation, these results demonstrate that irradiation conditions must be optimized in the context of the complete microfluidic system.

The optimized condition provided ∼85% cell viability at Day 1 while producing stable, monodisperse beads. Incomplete crosslinking can compromise bead stability and increase cellular exposure to reactive species^70,71^, while dissolved oxygen and unreacted methacrylate groups can further interfere with the photocrosslinking process^63,70^. Under the optimized conditions, the platform produced ∼200 µm sized microbeads with a CV of ∼5%, containing an average of ∼50 cells/bead, using different hydrogel combinations without the need of other modifications to the system. Thus, the system combines the bead dimensions required for multicellular microtissue formation with high cell occupancy, viability and production throughput.

This combination distinguishes our system from most previously reported high-throughput droplet and microgel platforms. Although several approaches achieve ∼liter/hour-scale production^15–17^, many generate droplets below 100 µm^16,17,72,73^, which are less suitable for multicellular encapsulation, and relatively few have demonstrated robust operation with highly concentrated cell-hydrogel suspensions. Step-emulsification further provides droplet sizes that are relatively insensitive to the flow-rate fluctuations, an important advantage for heterogeneous, cell-containing dispersed phases^73,74^. We have implemented this feature to facilitate upscaled production rates at the level up to ∼95 000 droplets/min.

In previous studies, translation of such high-throughput systems to cell encapsulation has been limited predominantly to single-cell encapsulation^75–81^. Reports on high-throughput multicellular encapsulation remain comparatively scarce. We have compared these previously reported systems to ours in terms of several performance metrics (Figure 7.a.). Headen *et al.* encapsulated 5–10 hMSCs per droplet but observed clogging above 5 million cells/mL cell loading concentration^82^, whereas Seeto *et al*. achieved much higher loading concentrations, they reported a rate of only ∼25 droplets/min^83^ (Figure 7.a.ii.). Other systems were limited to low loading concentrations or otherwise reported low number of cells per droplet^84,85^ or substantially lower throughput^84,86–89^ (Figure 7.a.i.). Zhang *et al*. reported number throughput reaching 700 000 droplets/min using 128 parallel flow-focusing junctions^90^, however, the cell loading concentration was limited to around 2 million/mL, which resulted typically in no more than a few (*N*∼5) cells/droplet. In-flow photocrosslinking of droplets has been demonstrated by Seeto *et al* and Mohamed *et al*, ^83,88^ but their volumetric flow rates were around 10 x lower than in our case. We directly checked that our D-chip is capable of generating droplets at the volumetric throughput even up to 300 uL/min, that is the highest volumetric throughput demonstrated so far in systems suited for multi-cell encapsulation. In summary, the key advantage of our system is the possibility of combining high volumetric throughput, droplet monodispersity and multicellular occupancy in a format directly compatible with microtissue engineering.

**Figure 7.**
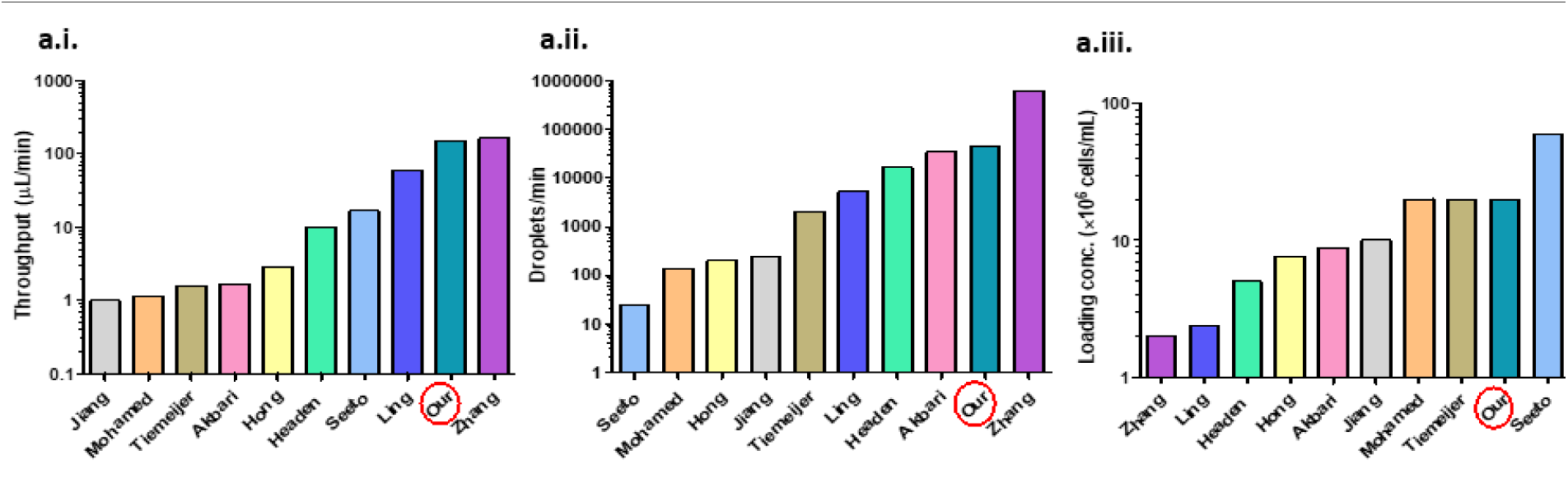
Comparative analysis with contemporary research. **a.** Microfluidic droplet generation for cell encapsulation applications compared to our system performance (in red oval) based on **i.** volumetric throughput, **ii.** droplet number throughput, and **iii.** loading concentration of cells (in million) for encapsulation.

We next demonstrated that different hydrogel compositions could be used with our high-throughput system for regulating microtissue formation. A formulation containing 4.8% GelMA, 0.4% HAMA, and 0.4% dECM (GelMA-HAMA-ECM) produced the fastest cell growth and microtissue formation. When the dECM-containing microdroplets were used for encapsulating MDA-MB-231 breast cancer cells, significantly enhanced cellular aggregation (approximately threefold larger than those in pure GelMA beads) was observed by the 5^th^ day of culture, accompanied by the highest metabolic activity. These results identify dECM-containing formulation as particularly promising for promoting rapid microtissue formation, consistent with the known ability of dECM to provide tissue-derived biochemical cues supporting proliferation and spheroid formation^91,92^. At the same time, the addition of HAMA served increased matrix stiffness and hydrogel stability^62,91^, without compromising the rapid MDA-MB-231 proliferation. Noteworthy, the impact of HAMA on cell proliferation rates, however, is not always positive and appears cell-type dependent. In our previous study, for example, we have found that HAMA inhibits leukemic and bone-marrow-derived cell models^93^.

Beyond disease modeling, the platform may be applicable to regenerative medicine. GelMA-HAMA-dECM hydrogels have recently been used for tailoring ADSC-laden scaffolds that promoted stem-cell differentiation in vivo^94^, suggesting that similar microbeads could serve as injectable cell-delivery constructs. More broadly, the scalability and tunable microenvironment of the present system could support the production of defined microtissues for disease modeling, drug testing, and regenerative applications.

## 5. Conclusion

In summary, we have developed a robust system for high-throughput encapsulation of large cell populations in hydrogel microbeads based on two connected chips and controlled photo-crosslinking. The system integrates the advantages of parallelization and step-emulsification, the latter warranting uninterrupted production of highly monodisperse microbeads despite fluctuations in cell concentration, hydrogel viscosity, etc. Homogeneous light exposure of each droplet, warranted by a dedicated crosslinking chip, as well as careful fine-tuning of UV wavelength and irradiance, altogether enable uniform polymerization, high cell viability, and long-term stability of the microbeads in culture. Additionally, we test the system using three different hydrogel compositions and determine that GelMA-HAMA-ECM is the most stable and valuable material for micro-tumor formation, e.g., in the case of triple negative breast cancer. We believe that the system can serve as a robust tool for microtissue modeling, with possible applications in organoid fabrication, high-throughput drug screening, and tissue/tumor phenotyping.

## Acknowledgements

We thank Patryk Adamczuk for assistance with micromilling and fabrication of the original microfluidic chips, Jhaleh Amirian for assistance with preliminary experiments during the initial development of the system, and Tomasz Dobrzański for fabrication of the HL lamp.

This work was funded by the NCN grant OPUS 17(2019/33/B/ST8/02145). The work of KG was financed by the National Centre for Research and Development under JTC 2022 Water4All Partnership (WATER4ALL/I/20/STARDUST/2024) program.

