## Supplementary Data for "High-throughput microfluidic encapsulation of cells in extracellular matrix microbeads for microtissue engineering"

### Slide 1
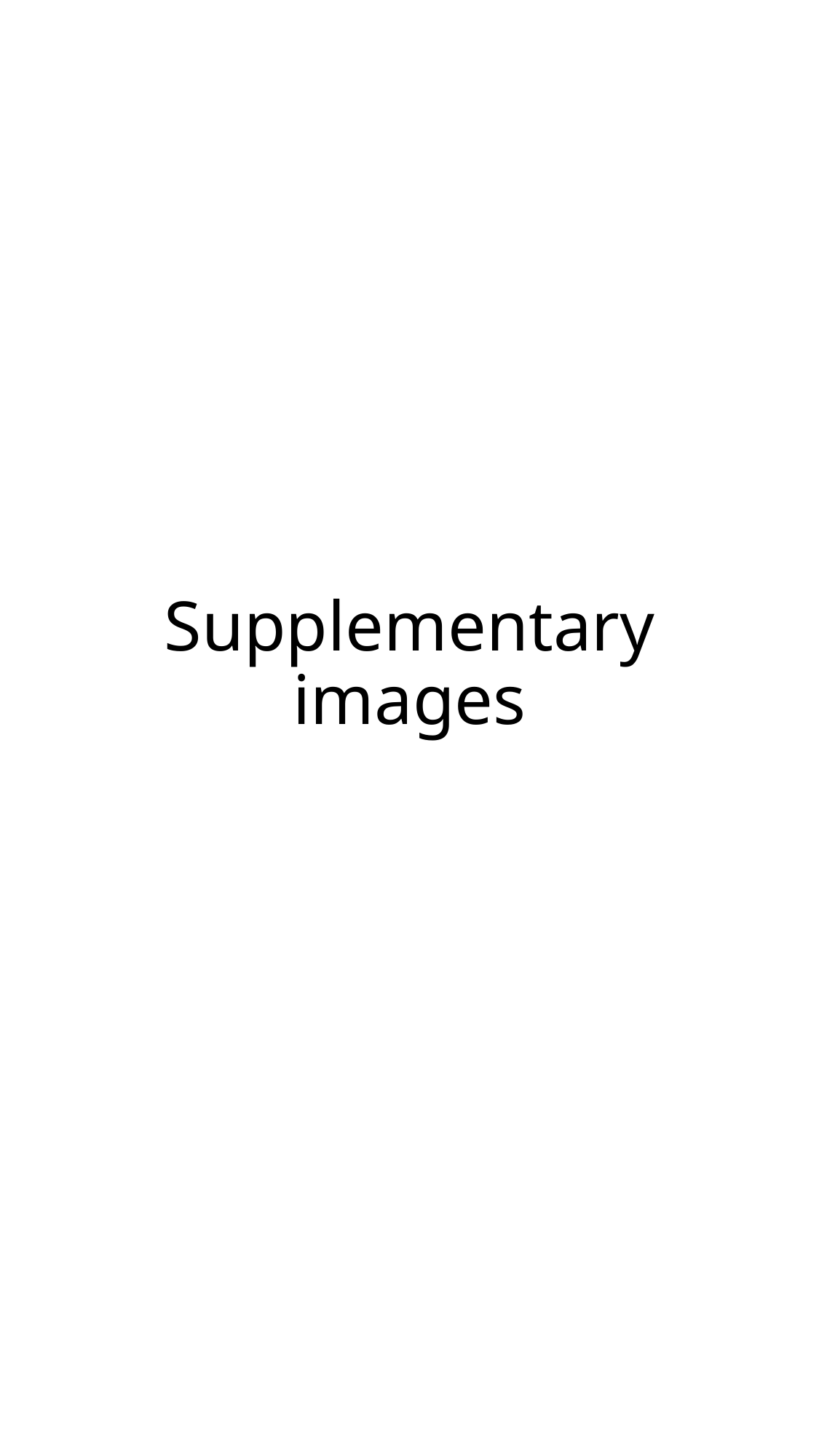

#### Supplementary images

### Slide 2
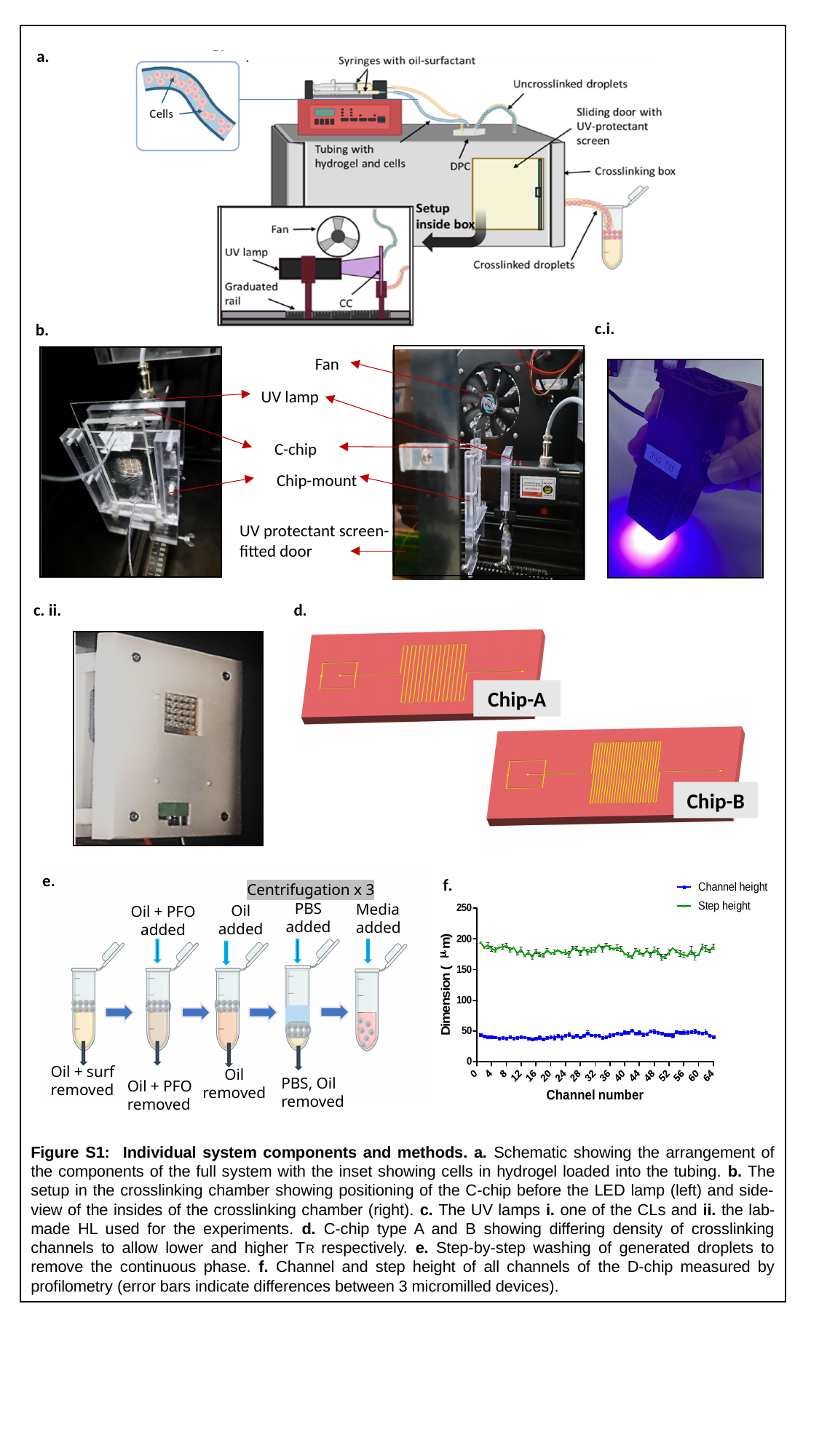

a.
d.ii.
c.i.
b.
Fan
UV lamp
C-chip
Chip-mount
UV protectant screen- fitted door
c. ii.
d.
Chip-A
Chip-B
Centrifugation x 3
PBS added
Media added
Oil added
Oil + PFO added
Oil + surf removed
Oil removed
PBS, Oil removed
Oil + PFO removed
e.
f.
Figure S1: Individual system components and methods. a. Schematic showing the arrangement of the components of the full system with the inset showing cells in hydrogel loaded into the tubing. b. The setup in the crosslinking chamber showing positioning of the C-chip before the LED lamp (left) and side-view of the insides of the crosslinking chamber (right). c. The UV lamps i. one of the CLs and ii. the lab-made HL used for the experiments. d. C-chip type A and B showing differing density of crosslinking channels to allow lower and higher TR respectively. e. Step-by-step washing of generated droplets to remove the continuous phase. f. Channel and step height of all channels of the D-chip measured by profilometry (error bars indicate differences between 3 micromilled devices).

### Slide 3
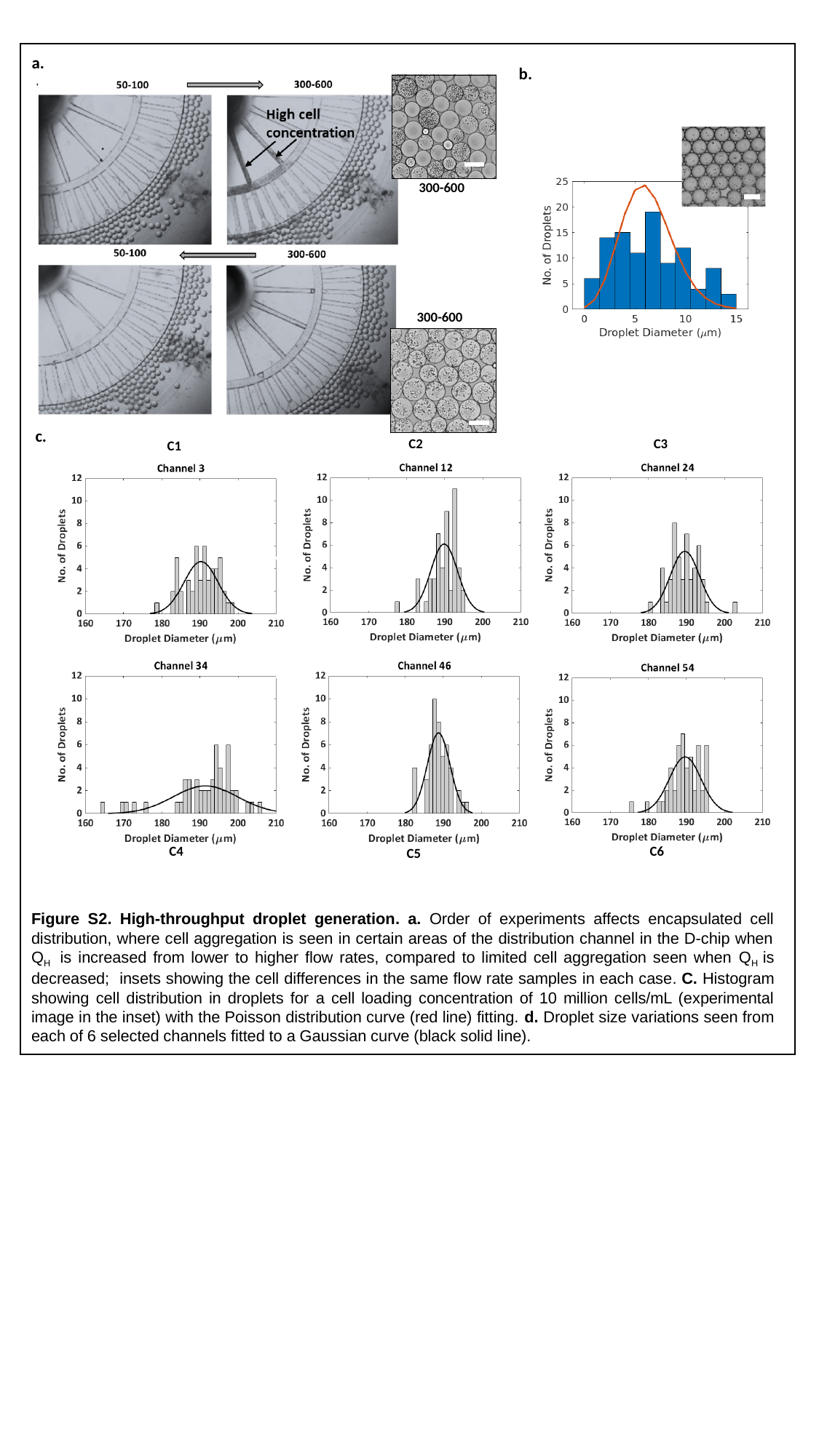

a.
b.
300-600
300-600
c.
C2
C3
C1
C6
C4
C5
Figure S2. High-throughput droplet generation. a. Order of experiments affects encapsulated cell distribution, where cell aggregation is seen in certain areas of the distribution channel in the D-chip when Q_H_ is increased from lower to higher flow rates, compared to limited cell aggregation seen when Q_H_ is decreased; insets showing the cell differences in the same flow rate samples in each case. C. Histogram showing cell distribution in droplets for a cell loading concentration of 10 million cells/mL (experimental image in the inset) with the Poisson distribution curve (red line) fitting. d. Droplet size variations seen from each of 6 selected channels fitted to a Gaussian curve (black solid line).

### Slide 4
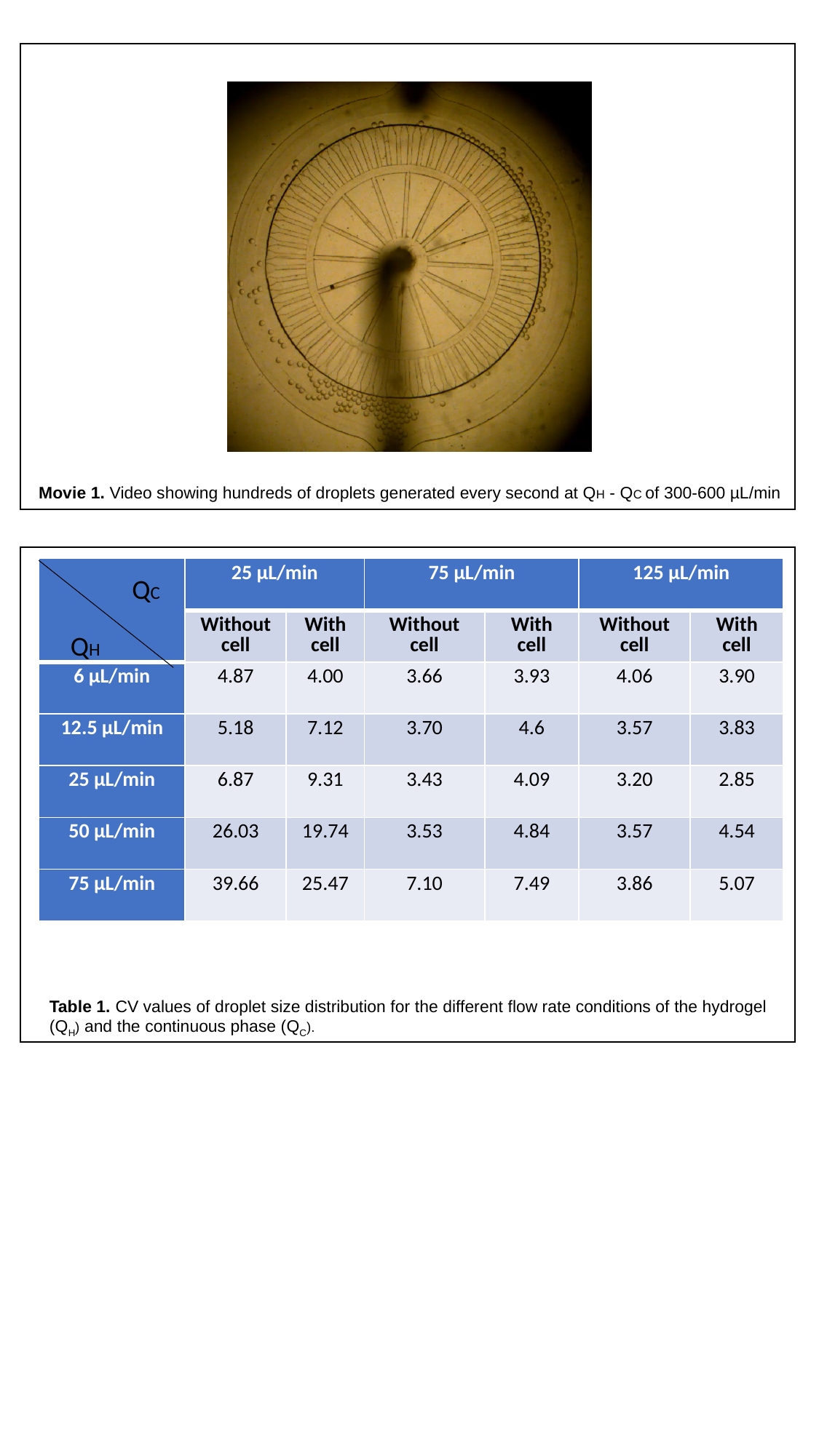

Movie 1. Video showing hundreds of droplets generated every second at QH - QC of 300-600 µL/min
| | 25 µL/min | | 75 µL/min | | 125 µL/min | |
| --- | --- | --- | --- | --- | --- | --- |
| | Without cell | With cell | Without cell | With cell | Without cell | With cell |
| 6 µL/min | 4.87 | 4.00 | 3.66 | 3.93 | 4.06 | 3.90 |
| 12.5 µL/min | 5.18 | 7.12 | 3.70 | 4.6 | 3.57 | 3.83 |
| 25 µL/min | 6.87 | 9.31 | 3.43 | 4.09 | 3.20 | 2.85 |
| 50 µL/min | 26.03 | 19.74 | 3.53 | 4.84 | 3.57 | 4.54 |
| 75 µL/min | 39.66 | 25.47 | 7.10 | 7.49 | 3.86 | 5.07 |
QC
QH
Table 1. CV values of droplet size distribution for the different flow rate conditions of the hydrogel (Q_H_) and the continuous phase (Q_C_).

### Slide 5
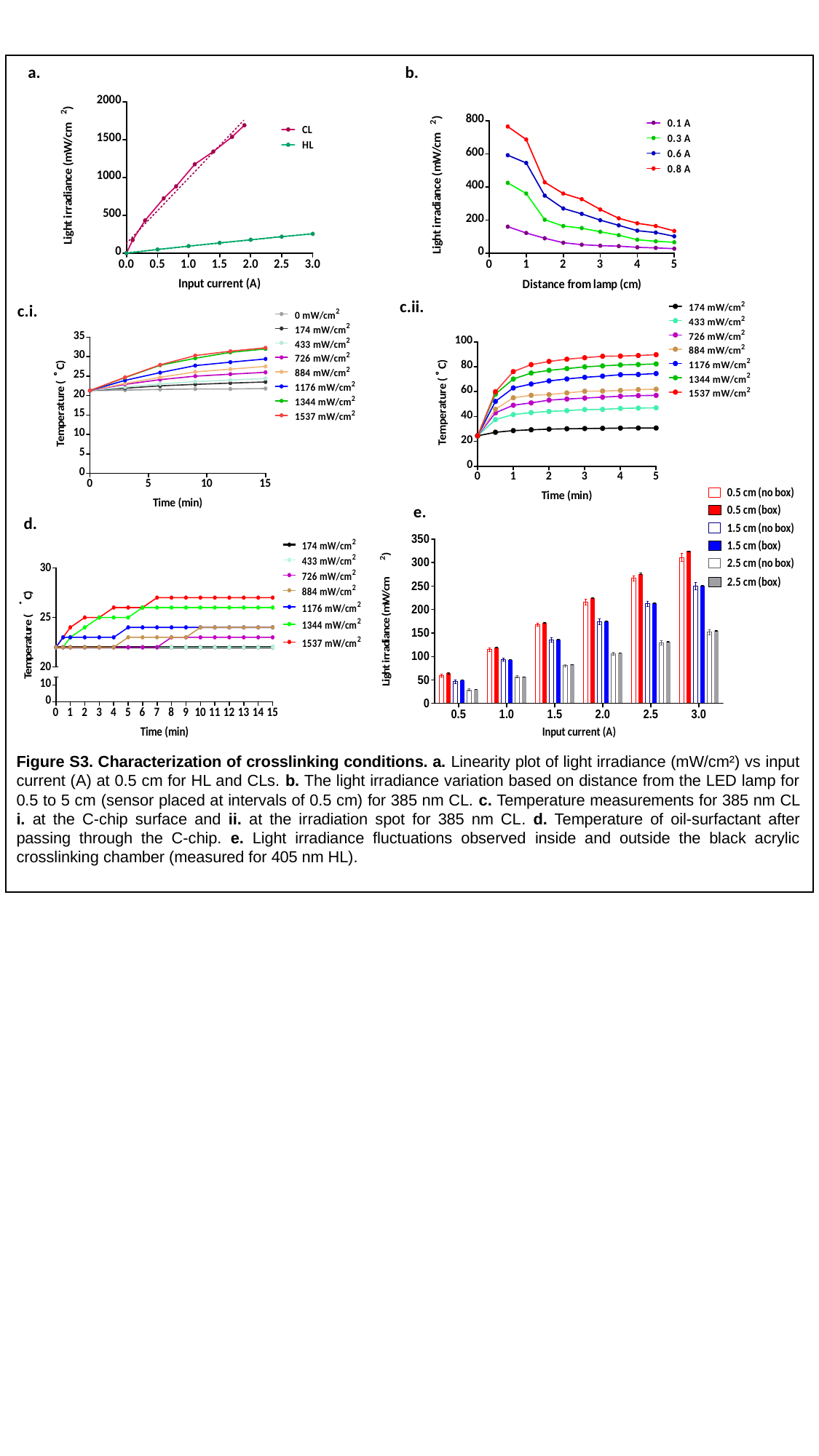

a.
b.
c.ii.
c.i.
e.
d.
Figure S3. Characterization of crosslinking conditions. a. Linearity plot of light irradiance (mW/cm²) vs input current (A) at 0.5 cm for HL and CLs. b. The light irradiance variation based on distance from the LED lamp for 0.5 to 5 cm (sensor placed at intervals of 0.5 cm) for 385 nm CL. c. Temperature measurements for 385 nm CL i. at the C-chip surface and ii. at the irradiation spot for 385 nm CL. d. Temperature of oil-surfactant after passing through the C-chip. e. Light irradiance fluctuations observed inside and outside the black acrylic crosslinking chamber (measured for 405 nm HL).

### Slide 6
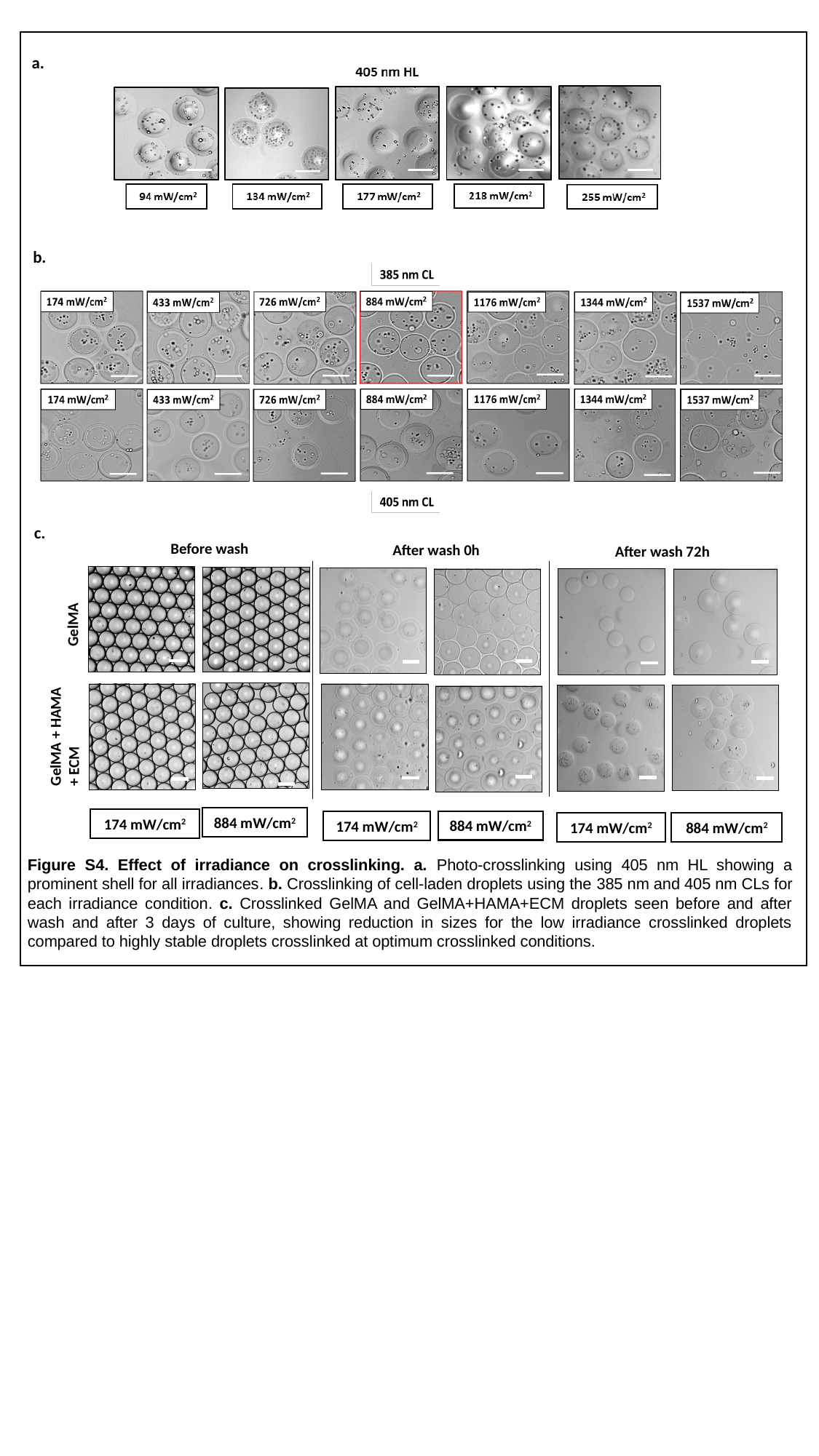

a.
b.
GelMA + HAMA + ECM
GelMA
Before wash
After wash 0h
After wash 72h
c.
884 mW/cm^2^
174 mW/cm^2^
884 mW/cm^2^
174 mW/cm^2^
884 mW/cm^2^
174 mW/cm^2^
Figure S4. Effect of irradiance on crosslinking. a. Photo-crosslinking using 405 nm HL showing a prominent shell for all irradiances. b. Crosslinking of cell-laden droplets using the 385 nm and 405 nm CLs for each irradiance condition. c. Crosslinked GelMA and GelMA+HAMA+ECM droplets seen before and after wash and after 3 days of culture, showing reduction in sizes for the low irradiance crosslinked droplets compared to highly stable droplets crosslinked at optimum crosslinked conditions.

### Slide 7
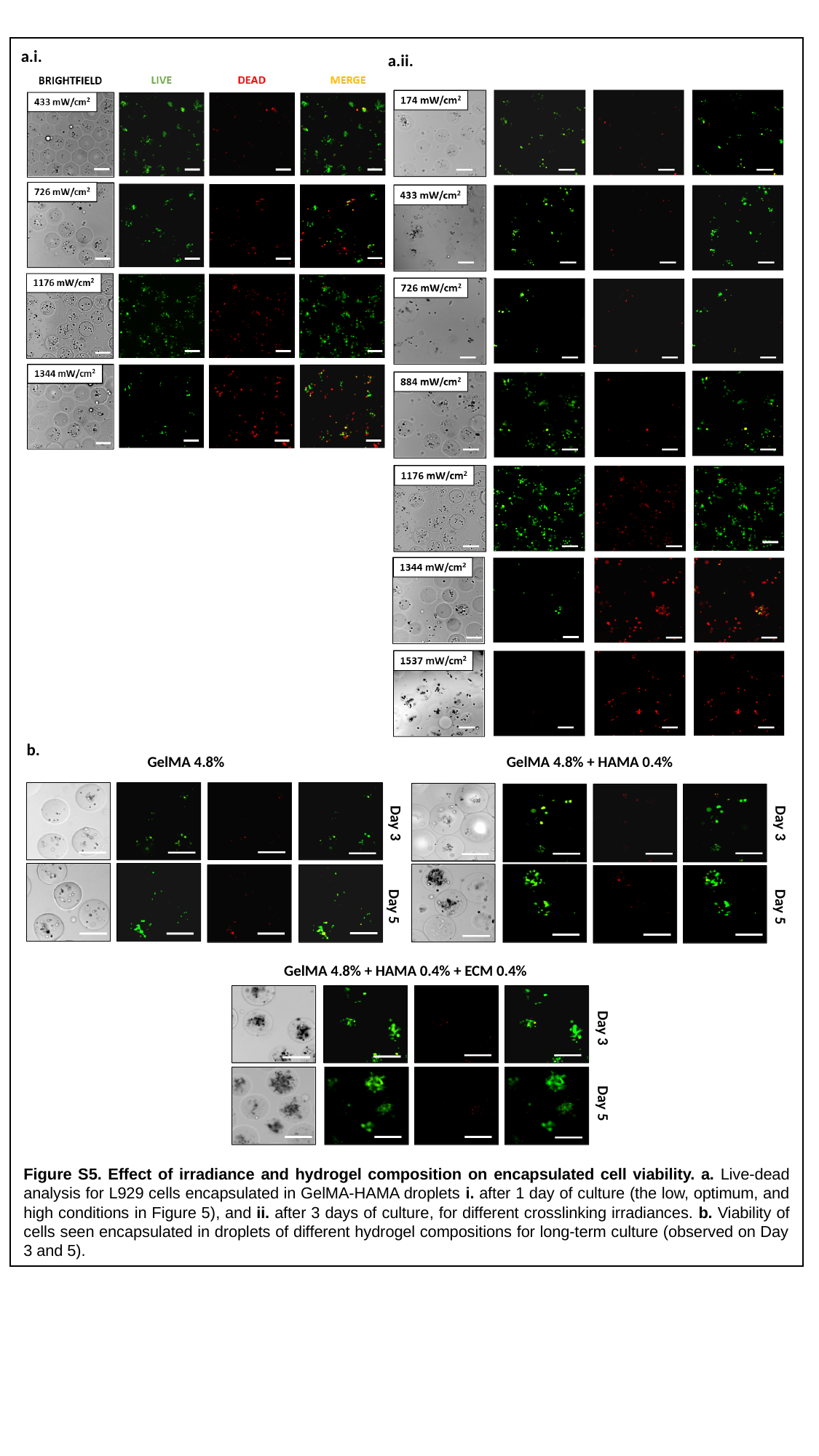

a.i.
a.ii.
b.
GelMA 4.8% + HAMA 0.4%
GelMA 4.8%
GelMA 4.8% + HAMA 0.4% + ECM 0.4%
Day 3
Day 3
Day 5
Day 5
Day 3
Day 5
Figure S5. Effect of irradiance and hydrogel composition on encapsulated cell viability. a. Live-dead analysis for L929 cells encapsulated in GelMA-HAMA droplets i. after 1 day of culture (the low, optimum, and high conditions in Figure 5), and ii. after 3 days of culture, for different crosslinking irradiances. b. Viability of cells seen encapsulated in droplets of different hydrogel compositions for long-term culture (observed on Day 3 and 5).
